# DNA replication is dispensable for developmental progression, but required for heterochromatin organization at mouse zygotic genome activation

**DOI:** 10.64898/2026.09.14.751383

**Authors:** Aurora Arroyo-Jimenez, Alicia Gallego, Antonio Barral, Maria Tiana, Elva Martin-Batista, Andrea Maidana, Miguel Manzanares, Marta Portela

**Author notes:** equal contribution (Aurora Arroyo-Jimenez, Alicia Gallego, Marta Portela).

## Abstract

Early mammalian development progresses through a small number of cleavage divisions with extensive transcriptional and epigenetic reprogramming that result in the very first lineage decisions, leading to blastocyst formation. Although classical embryological studies suggested that developmental progression can occur independently of normal cell division, the extent to which DNA replication contributes to these processes remains unclear. In particular, whether replication is required simply for proliferation or also for establishing the cellular states that regulate stage-specific gene expression is unknown. Here we show that mouse preimplantation development can proceed without DNA replication, but it is required to maintain proper heterochromatin organization at the 2-cell stage, the time of zygotic genome activation. The inhibition of DNA replication with aphidicolin blocked cleavage divisions without preventing zygotic genome activation, embryo morphogenesis, activation of lineage-specification factors, or the establishment of stage-specific transcriptional programs. However, replication arrest selectively altered the expression of developmental genes associated with late-replicating, Polycomb-enriched chromatin domains, and also reduced global levels of H3K27me3 and H3K9me3. These findings indicate that developmental timing in the early embryo is largely uncoupled from DNA replication and cell-cycle progression. Instead, DNA replication contributes to transcriptional fidelity by establishing or maintaining repressive chromatin states during the extensive genome reprogramming that accompanies zygotic genome activation.

## INTRODUCTION

Following fertilization, mammalian embryos undergo relatively slow cleavage divisions, while simultaneously experiencing extensive transcriptional and epigenetic reprogramming. These coordinated events culminate in the formation of a ∼100-cell blastocyst stage embryo, composed of three distinct lineages, and ready for implantation in the maternal uterus (Fig. 1).

**Figure 1.**
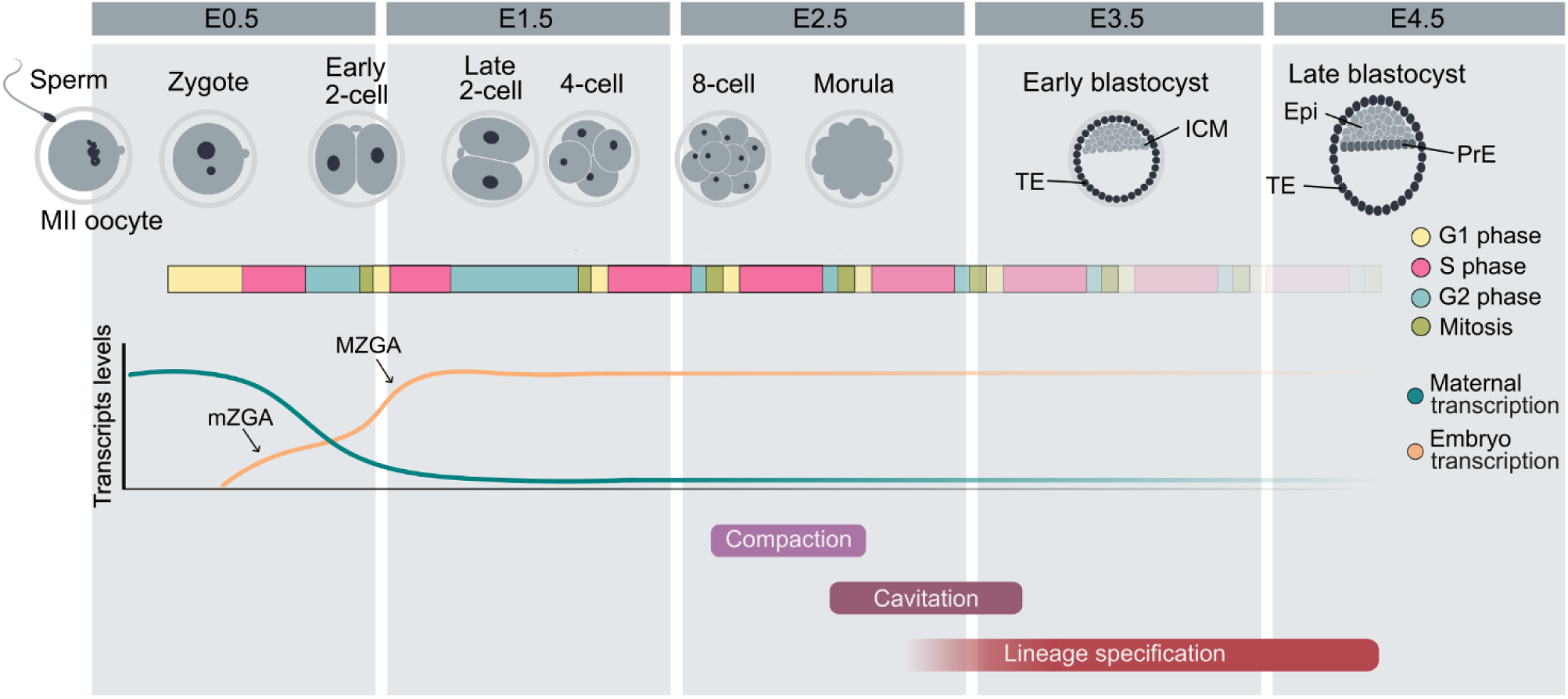
Developmental and transcriptional events during mouse preimplantation embryogenesis. Diagram summarizing mouse preimplantation development from fertilization (E0.5) to the late blastocyst stage (E4.5). Developmental progression, cell-cycle phases (colour-coded), and transcription dynamics are shown. Maternal transcripts progressively decline, while embryonic transcription is initiated during minor zygotic genome activation (mZGA) and reaches full activation during major zygotic genome activation (MZGA) at the 2-cell stage. Compaction, cavitation, and lineage specification occur sequentially as embryos progress through the morula and blastocyst stages, giving rise to the trophectoderm (TE), inner cell mass (ICM), epiblast (Epi), and primitive endoderm (PrE).

Although these processes are temporally coordinated, classical embryological studies have revealed that certain developmental transitions, including zygotic genome activation and blastocyst formation, can proceed independently of normal cleavage progression (Newport and Kirschner, 1982; Pratt et al., 1981). Even under cell cycle arrest, embryos from diverse species are able to proceed with morphogenesis and correctly express markers of differentiated cell types (Edgar and O’Farrell, 1989; Rollins and Andrews, 1991; Nair et al., 2013). These observations suggest that early developmental progression is not strictly governed by the number of cleavages, but instead follows intrinsic temporal programs (Dean and Rossant, 1984). This evidence, together with the technical challenges of studying cell cycle regulation in embryos composed of only a few cells, has shifted attention away from replication progression as a potential driver of developmental progression. As a result, the extent to which DNA replication itself contributes to embryonic timing and developmental transitions remains largely unresolved.

However, the first embryonic cell cycles are not merely proliferative events. In the mouse, they take place together with extensive chromatin reorganization, including global epigenetic reprogramming, progressive establishment of replication timing (RT) domains, and the consolidation of higher-order genome organization (Eckersley-Maslin et al., 2018; Collombet et al., 2020; Nakatani et al., 2024; Xu et al., 2024). Although S-phase duration remains relatively constant across early cleavages (Fig. 1) (Ciemerych et al., 1999, Artus and Cohen-Tannoudji, 2008), replication dynamics change substantially, with slow fork progression in early stages and marked acceleration toward the blastocyst stage (Ahuja et al., 2016; Nakatani et al., 2022). Furthermore, inhibition of DNA replication interferes with maternal DNA demethylation and the restoration of three-dimensional chromatin architecture (Ke et al., 2017; Eckersley-Maslin et al., 2018), indicating that replication is intimately linked to genome organization.

These findings raise a fundamental question: is DNA replication in the early embryo simply related to cell proliferation, or does it play a specific role in establishing the dynamic chromatin states that underlie stage-specific gene regulation? In this study, we address this question by examining the consequences of transient replication inhibition during mouse preimplantation development, focusing on how perturbation of S-phase impacts lineage specification and the proper implementation of developmentally regulated transcriptional programs.

## RESULTS

### Inhibition of DNA replication in preimplantation embryos

To dissect the role of DNA replication during mouse preimplantation development, we disrupted cell-cycle progression by treating embryos with aphidicolin during the second and third cleavage cycles (Fig. 1). These stages encompass key developmental transitions: zygotic genome activation (ZGA), that occurs in two phases, minor ZGA from the zygote to the early 2-cell (E2C), followed by major ZGA at the transition from E2C to late 2-cell (L2C) stage; and the establishment of apico-basal polarity at the 4-cell (4C) to 8-cell (8C) transition, preceding compaction and the first lineage segregation into trophectoderm (TE) and inner cell mass (ICM) (Chazaud and Yamanaka, 2016). Aphidicolin was selected as a specific and reversible inhibitor of replicative DNA polymerases, enabling transient block of DNA synthesis at S-phase and preventing cell-cycle progression (Ikegami et al., 1978; Spadari et al., 1982), that has been previously used to block replication during mouse preimplantation development (Bolton et al., 1984; Dean and Rossant, 1984; Smith and Johnson, 1985; Spindle et al., 1985).

We first checked that DNA replication was effectively inhibited by treating E2C embryos with aphidicolin for 16 h. Treatment was initiated immediately after completion of the first cell division, ensuring that S phase had not yet begun, and continued until the L2C stage (Fig. 2A). DNA synthesis was monitored by incorporation of EdU (5-ethynyl-2′-deoxyuridine), a synthetic thymidine analog, that was added together with aphidicolin (Fig. 2A). EdU incorporation was drastically reduced in aphidicolin-treated embryos (Fig. 2B), indicating that DNA replication was inhibited almost immediately after adding aphidicolin. Interestingly, nuclear volume or nuclear morphology did not change in treated embryos (Fig. S1A).

**Figure 2.**
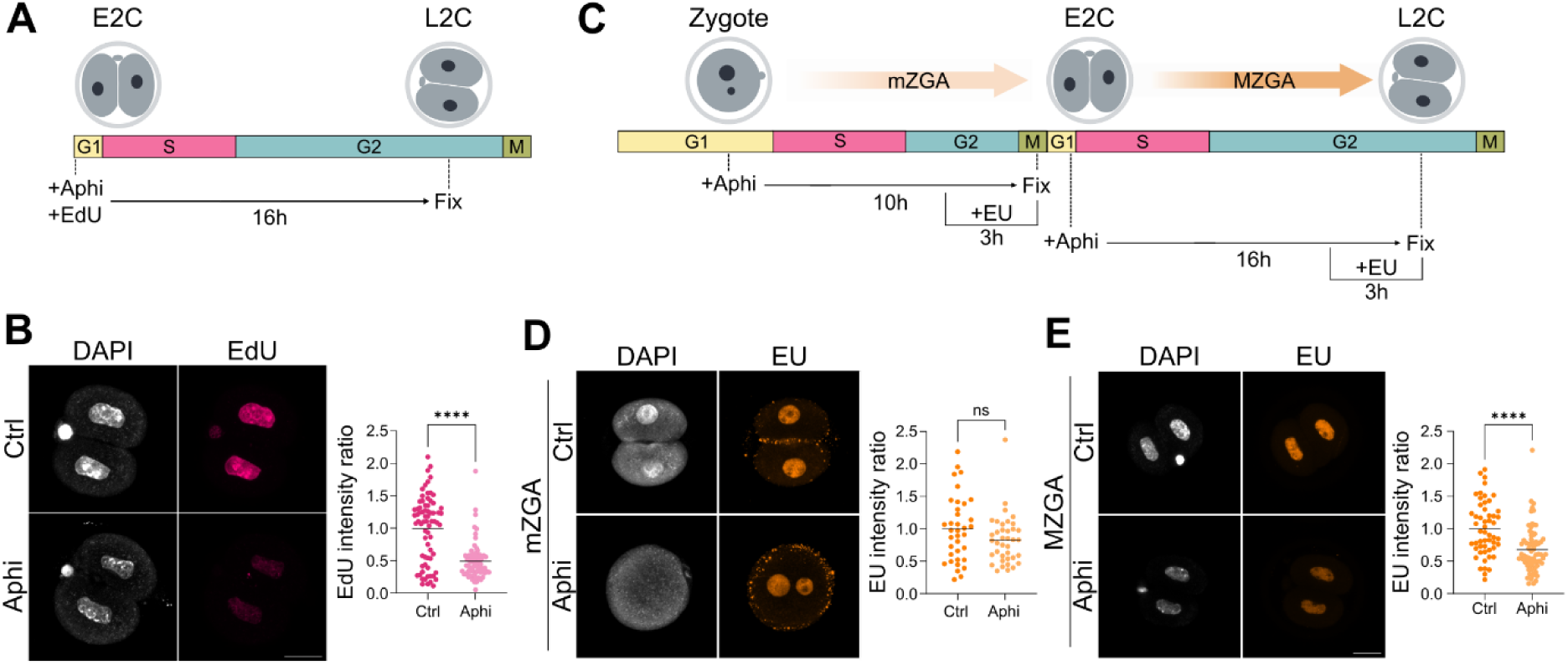
DNA replication and transcriptional dynamics at zygotic genome activation in mouse embryos. **(A)** Schematic of cell-cycle progression from early 2-cell (E2C) to late 2-cell (L2C) stage mouse embryos and experimental design to check DNA synthesis upon aphidicolin treatment. Cell-cycle phases (G1, S, G2, and M) and windows of treatment are indicated below. **(B)** Representative images of L2C control and age-matched aphidicolin-treated embryos labeled for DNA synthesis (EdU, magenta) and total DNA (DAPI, gray). Quantification of EdU signal is shown on the right, as the ratio between the mean intensity of the nuclei from each embryo and the mean intensity of control embryos. Each dot represents an individual embryo (control, n=72; Aphi, n=66), and horizontal lines indicate the mean. **** P<0.001; two-tailed Student’s t-test. **(C)** Schematic of cell-cycle progression from one-cell (Zygote) to L2C stage mouse embryos and experimental design to check transcription at zygotic genome activation upon aphidicolin treatment. Cell-cycle phases (G1, S, G2, and M) and windows of treatment are indicated below. Arrows denote the temporal windows of minor (mZGA) and mayor (MZGA) zygotic genome activation. **(D)** Representative images of E2C control and age-matched aphidicolin-treated embryos labeled for RNA synthesis (EU, orange) and DNA (DAPI, gray) at the mZGA. Aphidicolin-treated embryo is at the one-cell stage and shows two unfused pronuclei. Quantification of EU fluorescence intensity is shown on the right, as the ratio between the mean intensity of the nuclei from each embryo and the mean intensity of control embryos. Each dot represents an individual embryo (control, n=36; Aphi, n=38), and horizontal lines indicate the mean. ns, non-significant; two-tailed Student’s t-test. **(E)** Representative images of L2C control and age-matched aphidicolin-treated embryos labeled for RNA synthesis (EU, orange) and DNA (DAPI, gray) at the MZGA. Quantification of EU fluorescence intensity is shown on the right, as the ratio between the mean intensity of the nuclei from each embryo and the mean intensity of control embryos. Each dot represents an individual nucleus (control, n=53; Aphi, n=71), and horizontal lines indicate the mean. **** P<0.001; two-tailed Student’s t-test.

### Zygotic genome activation occurs in the absence of DNA replication

Next, we investigated how blocking replication affects the initiation of zygotic transcription in the embryo. To assess minor ZGA, one-cell zygotes were treated with aphidicolin for 10 h, by which time control embryos have completed the first cell division and reached the E2C stage (Fig. 2C). General transcriptional activity was measured by incorporation of EU (5-ethynyluridine), a synthetic uridine analog, into nascent RNA. EU was added 3 h before embryo fixation. Blocking the first round of DNA replication led to arrest at the one-cell stage, as has been previously described (Bolton et al., 1984; Xu et al., 2024), with the vast majority ofembryos still retaining two separate pronuclei (Figs 2D, S1B). Despite this, treated embryos showed transcription levels comparable to controls (Fig. 2D). To study major ZGA, embryos were treated with aphidicolin from the E2C to the L2C stage for 16 h with a final 3h EU treatment before fixation (Fig. 2C). In this case, we observed a small but significant decrease in EU incorporation (Fig. 2E), implying that inhibition of the second round of DNA replication affects de novo zygotic transcription.

### Morphogenesis and lineage specification are not dependent on DNA replication

To further explore the role of DNA replication in mouse early embryos, we blocked the third cleavage, that takes place between 4C and 8C stage embryos (Fig. 1). At this developmental window, key morphological events are triggered, such as blastomere polarization and embryo compaction, processes that culminate at the morula stage (Chazaud and Yamanaka, 2016). We observed that aphidicolin treated embryos at the 4C stage for 16 h (Fig. 3A) did not divide, but showed morphological evidence of compaction between blastomeres similar to that observed in age-matched 8C controls (Fig. 3B), consistent with earlier reports (Smith and Johnson, 1985). To assess the apico-basal polarization of blastomeres, which occurs concomitantly with compaction (Ziomek and Johnson, 1980), we analyzed the distribution of phosphorylated EZRIN (pEZR), a marker that accumulates at the apical, outward-facing surface of blastomeres at the 8C stage (Louvet et al., 1996). Aphidicolin-treated embryos, although remaining at 4 cells, displayed apical enrichment of pEZR similar to that observed in 8C embryos and absent from control 4C embryos (Fig. 3C). Morphometric analysis of embryos revealed that total cellular volume and surface area of aphidicolin-treated embryos were more similar to those of 4C controls than to 8C age-matched controls. However, nuclear volume was markedly increased compared to both 4- and 8C controls, resulting in a nuclear-to-cytoplasmatic (N/C) ratio of aphidicolin-treated embryos similar to 8C embryos (Fig. S1C). Overall, these results indicate that DNA replication and the progression from 4 to 8 cells are dispensable for the very first morphological changes that take place during mouse preimplantation development.

**Figure 3.**
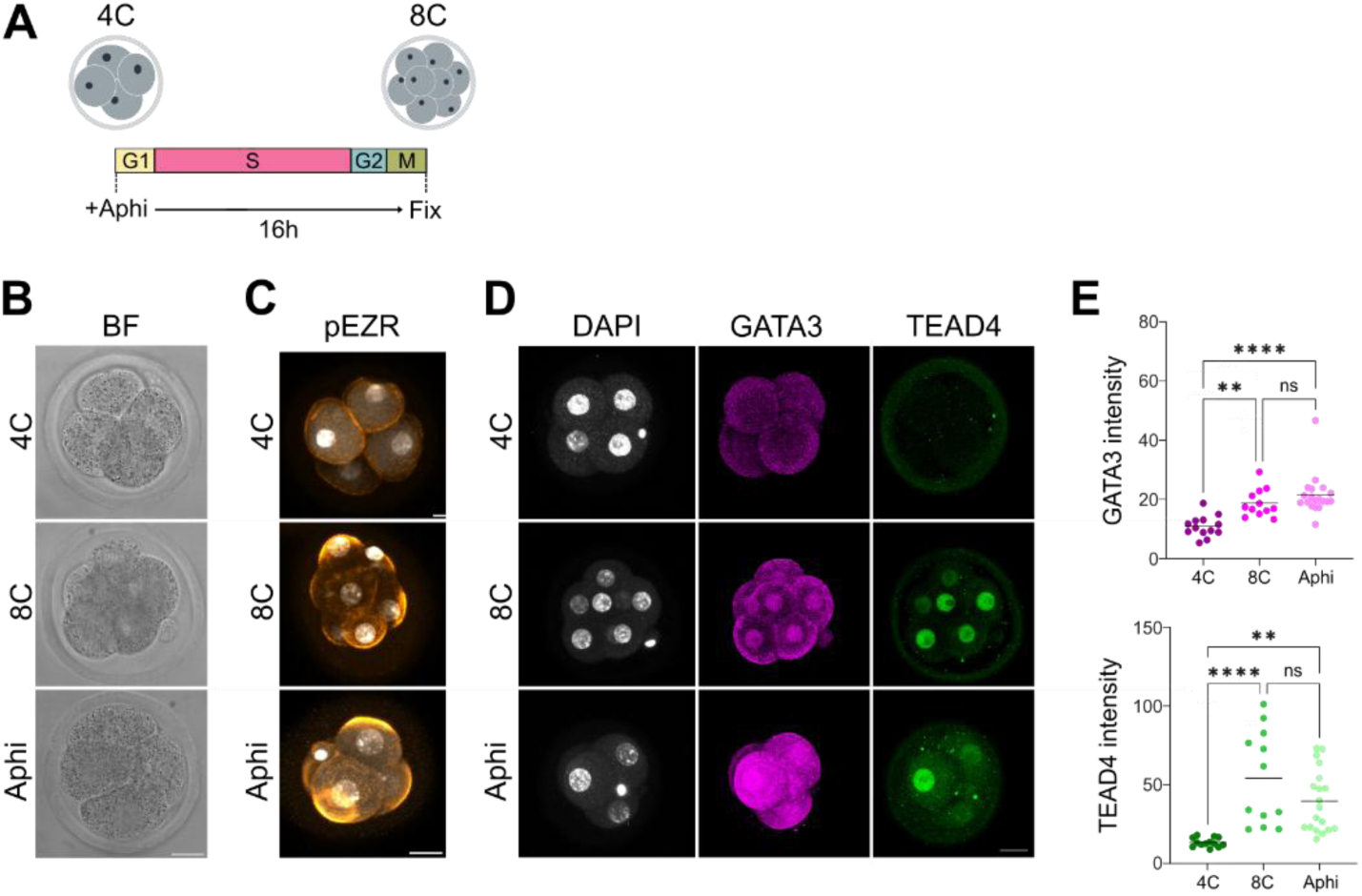
Early lineage specification in the absence of DNA replication. **(A)** Schematic of cell-cycle progression from 4-cell (4C) to 8-cell (8C stage mouse embryos. Cell-cycle phases (G1, S, G2, and M) and the window of aphidicolin treatment is indicated below. **(B)** Representative bright-field (BF) images of 4C, 8C and age-matched aphidicolin-treated embryos. Blastomere compaction is observed in both 8C and aphidicolin-treated embryos. **(C)** Representative images of 4C, 8C and age-matched aphidicolin-treated embryos labeled with an antibody that recognizes phospho-EZRIN (pEZR, orange) and with DAPI to mark the nucleus (gray). **(D)** Representative images of 4C, 8C and age-matched aphidicolin-treated embryos labeled with antibodies for GATA3 (middle, magenta) or TEAD4 (left, green). Nuclei are stained with DAPI (right, gray). **(E)** Quantification of the fluorescent intensity in 4C, 8C and age-matched aphidicolin-treated embryos, labeled with antibodies that recognize GATA3 (left, magenta) or TEAD4 (right, green). Each dot represents an individual embryo (4C, n=13; 8C, n=12; Aphi, n=19). ** P<0.01; **** P<0.001; ns, non-significant; two-tailed Student’s t-test.

To investigate the impact of cell cycle arrest on lineage decisions, we examined the activation of cell-fate associated transcription factors at the 4C to 8C transition, as the 8C stage marks the onset of expression of key factors involved in TE specification, including GATA3 or TEAD4 (Nishioka et al., 2008; Home et al., 2009). As expected, neither factor was detected in 4C embryos, whereas both were robustly upregulated in 8C embryos. Remarkably, aphidicolin-treated embryos exhibited levels of TEAD4 and GATA3 comparable to those observed in 8C embryos despite remaining at 4 cells (Figs. 3D, E). These findings indicate that replication inhibition does not disrupt early genetic programs involved in lineage decisions. Rather, coordinated morphogenetic processes continue to unfold with appropriate developmental timing, independently of genome replication, cell division, and cell number.

### Global transcription is largely maintained after DNA replication arrest

To gain further insight into more precise effects of the transcriptional response to aphidicolin block during preimplantation development, we performed RNA sequencing (RNA-seq) at both developmental time windows described above, spanning the second and third rounds of DNA replication: from E2C to L2C (Fig. 2A), and from the 4C to the 8C stage (Fig. 3A). For each window, pools of 20-60 embryos were collected and sequenced, including control embryos at the initial and final time points, plus age-matched aphidicolin-treated embryos at the final time point.

Principal component analysis (Fig. 4A, B) and hierarchical clustering (Fig. S2A, B) revealed that, in both time windows, aphidicolin-treated embryos clustered more closely with their chronologically matched controls (L2-cell, 8C) than with replication-stage matched controls (E2-cell, 4C), indicating that transcriptional progression was more strongly associated with developmental time than with replication-cycle progression.

**Figure 4.**
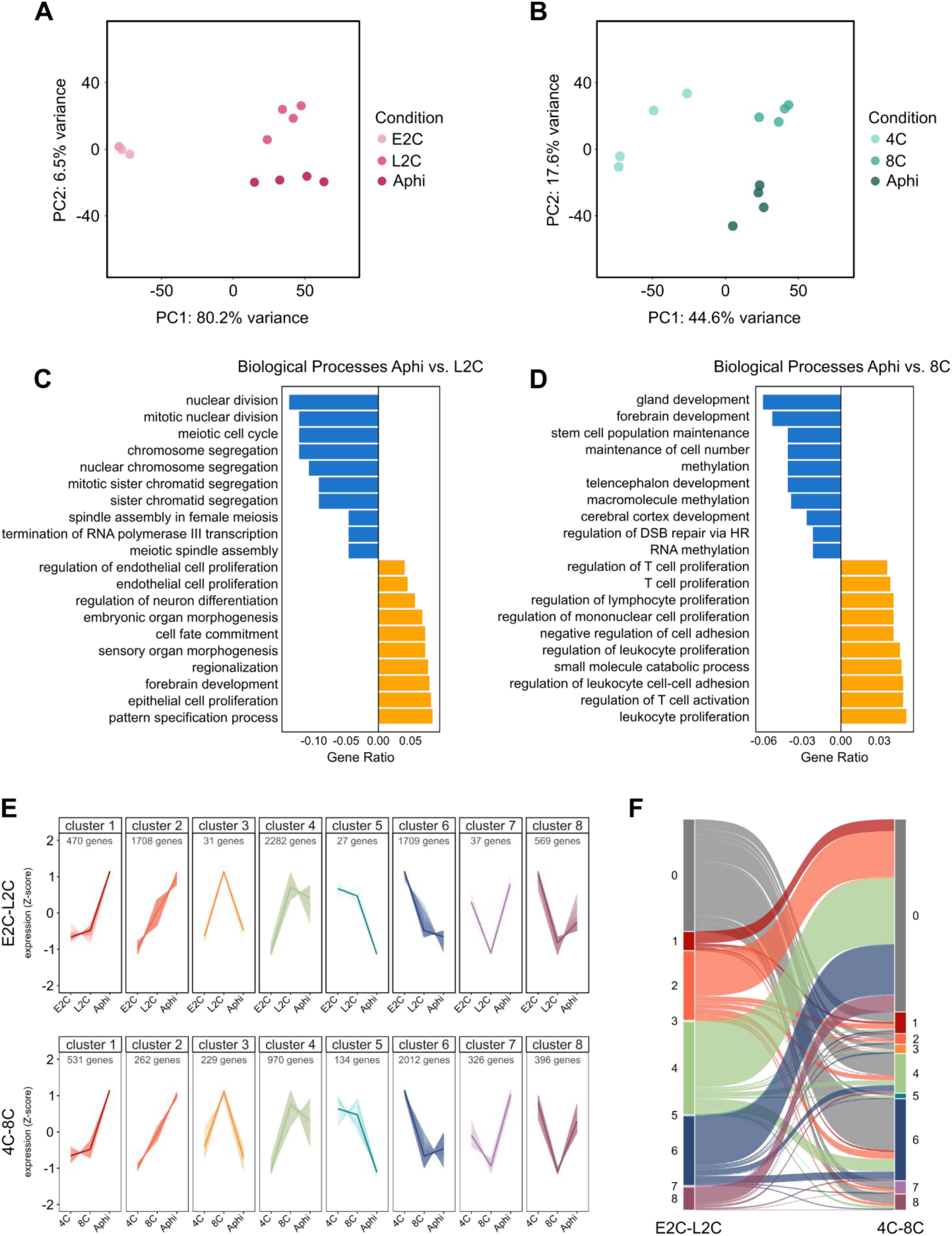
Transcriptional response to DNA replication block in preimplantation embryos. **(A)** Principal component analysis of RNA-seq transcriptomic profiles from E2C control, L2C control, and age-matched aphidicolin-treated (Aphi) embryos. Each point represents one biological replicate (n=4 per group). PC1 and PC2 explain 80.2% and 6.5% of the variance, respectively. **(B)** Principal component analysis of RNA-seq transcriptomic profiles from 4C control, 8C control, and age-matched aphidicolin-treated (Aphi) embryos. Each point represents one biological replicate (n=4 per group). PC1 and PC2 explain 44.6% and 17.6% of the variance, respectively. **(C)** GO Biological Process enrichment analysis of up (orange) and down (blue) DEGs between aphidicolin-treated embryos at the E2C stage for 16 h and L2C control embryos. Top ten statistically significant categories (adj P-value<0.01) are shown, ranked by gene ratio (Table S2). **(D)** GO Biological Process enrichment analysis of up (orange) and down (blue) DEGs between aphidicolin-treated embryos at the 4C stage for 16 h and 8C control embryos. Top ten statistically significant categories (adj P-value<0.01) are shown, ranked by gene ratio (Table S2). **(E)** Clustering analysis of differentially expressed genes (DEGs) identified in the E2C-to-L2C transition (top; 6,833 genes) and the 4C-to-8C transition (bottom; 4,860 genes). Genes were grouped into eight clusters according to their expression dynamics across developmental stages and age-matched aphidicolin-treated embryos (Aphi). Expression levels are shown as z-scores, and the number of genes in each cluster is indicated. The x-axis represents E2C, L2C, and Aphi samples (top) or 4C, 8C, and Aphi samples (bottom). Lines represent the average expression pattern of genes within each cluster, and shaded regions indicate variability within the cluster. **(F)** Alluvial plot showing the correspondence between gene clusters identified in the E2C-to-L2C dataset (left) and the 4C-to-8C dataset (right). Each ribbon represents genes shared between clusters in the two analyses, and ribbon width is proportional to the number of genes. Colors indicate the cluster assignment in the E2C-to-L2C dataset. The plot highlights the extent to which gene expression patterns are conserved or redistributed between the two developmental transitions.

Differential gene expression analysis (Table S1) revealed that transcriptomic changes were more pronounced during the E2C-to-L2C transition than in the 4C to 8C window. Approximately two-thirds of differentially expressed genes (DEGs) between L2C and E2C embryos were upregulated, reflecting the strong transcriptional stimulation at the ZGA (Fig. S2C). Also, most of DEGs at E2C vs L2C also changed in the aphidicolin vs E2C comparison (Fig. S2D). However, only 555 genes differed between aphidicolin-treated and L2C embryos (Fig. S2E), of which 87% were upregulated. In contrast, when comparing 8C and 4C embryos, the majority of DEGs were downregulated (Fig. S2F), and also differentially expressed in the aphidicolin vs 4C comparison (Fig. S2G). Finally, 1,322 DEGs distinguished aphidicolin-treated from 8C embryos (Fig. S2H), approximately two-thirds of them being upregulated.

One possibility we could not rule out is that aphidicolin treatment led to a general transcriptional response to its known deleterious effects, including replication stress, DNA damage and apoptosis (Zeman and Cimprich, 2014). To check so, we examined genes differentially expressed by aphidicolin treatment across all four stages analyzed, and found that only 43 were shared (Fig. S2I), indicating the absence of a broad and uniform transcriptional effect of aphidicolin treatment. A subset of these commonly upregulated genes (*Btg2*, *Cdkn1c*, *Ccng1*, *Fosl1*, *Phlda3, Plk2, Sapcd2*) is consistent with a p53-mediated stress response and growth arrest, which represent canonical cellular responses to aphidicolin-induced replication stress. This conclusion was further supported by comparison with a published dataset of transcriptional response of mouse embryonic fibroblasts (MEF) in response to aphidicolin (Mazouzi et al. 2016). When comparing changes induced by aphidicolin in L2C or 8C embryos with those observed in MEFs after 24 h of treatment, only 25 genes were shared (Fig. S2J). Despite this limited overlap, the shared genes included several associated with cell cycle (*Ccnd2*, *Ccng1*), immediate early genes (*Atf3*, *Fos*, *Fosl1*, *Junb*) and stress response (*Btg2*, *Phlda3*, *Plk2*).

Gene Ontology (GO) analysis of DEGs also highlighted stage-specific responses. When E2C embryos were compared with L2C embryos, genes upregulated in E2C embryos were enriched for categories related to DNA dynamics and cell-cycle processes, whereas genes upregulated in L2C embryos were related to ribosome biogenesis and translation (Fig. S3A, Table S2). This observation is consistent with E2C embryos being actively engaged in the first embryonic cell divisions, while L2C embryos are preparing for the massive de novo translation associated with ZGA, and match recent high-resolution analysis of transcriptional changes at ZGA (Al-Mousawi et al., 2026). Similarly, genes upregulated in 4C embryos were enriched in terms related to regionalization and developmental processes, whereas genes upregulated in 8C embryos were enriched in terms for metabolic processes (Fig. S3B, Table S2). As expected, comparisons with aphidicolin-treated embryos showed GO enrichments closely resembling normal developmental progression (Fig. S3A, B; Table S2).

The comparison of aphidicolin-treated embryos to stage-matched control was more informative. Genes whose expression was reduced by aphidicolin were related to cell division and cell-cycle progression in embryos treated from E2C to L2C embryos, and to DNA repair, maintenance of cell number and development in embryos treated from 4C to 8C. On the other hand, genes upregulated by aphidicolin in L2C embryos showed enrichment of developmental GO terms, particularly those related to neural development, whereas in 8C embryos these were predominantly enriched in categories related to control of cell proliferation and metabolic processes (Fig. 4C, D; Table S2).

To determine whether our data reflected appropriate temporal progression, we compared our dataset with previously defined potency-associated gene sets (Yang et al., 2022). Embryos in the E2C-to-L2C time window properly activated minor ZGA and major ZGA genes, whereas embryos in the 4C-to-8C window repressed totipotency-associated genes while activating pluripotency-associated genes (Fig. S3C). In line with the results described above (Figs 4A, 4B, S2A, S2B), aphidicolin-treated embryos behaved similarly to L2C or 8C control embryos.

### Selective effects of DNA replication on developmental gene expression trajectories

To better understand the transcriptional changes occurring in embryos when DNA replication is blocked, we carried out a clustering analysis for both developmental windows. To this end, we selected all genes that were differentially expressed in at least one of the three pairwise comparisons for each window (E2C vs L2C, E2C vs aphidicolin-treated embryos, and L2C vs aphidicolin-treated embryos; and equally for the 4C-to-8C window; Table S1). Genes from each window were assigned to one of eight clusters representing different trajectories (Fig. 4E, Table S1). A ninth cluster (cluster #0), used in subsequent analysis as a control group, included genes that were expressed but did not change in any of the comparisons. In agreement with the initial analysis (Fig. 4A, B), the majority of genes were unaffected by aphidicolin treatment, and belonged to cluster #4 (genes that increased expression along developmental progression and remained at similar levels in aphidicolin-treated embryos) and cluster #6 (genes whose expression decreased as development progresses and were unchanged by aphidicolin). Together, these two clusters accounted for 58.4% of deregulated genes in the E2C-to-L2C dataset, and 61.4% of the genes in the 4C-to 8C dataset. The remaining clusters represent different responses to aphidicolin, such as clusters #1 and #2, which include genes whose expression is markedly upregulated upon aphidicolin treatment. It should be noted that clusters #3, #5 and #7 contained very few genes in the E2C-to-L2C dataset, limiting the robustness of downstream analyses using data from these clusters.

Given the similar expression trajectories observed in both datasets, we investigated whether aphidicolin affected similar subsets of genes across the two developmental windows. To do so, genes from clusters #0 to #8 in the E2C-to-L2C dataset were mapped onto clusters in the 4C-to-8C dataset. If some clusters represented a conserved aphidicolin-responsive gene set acting independently of developmental stage, a strong one-to-one correspondence between clusters would be expected. However, we observed an extensive redistribution of genes across multiple clusters, with no clear cluster-to-cluster correspondence between the two datasets (Fig. 4F). These results indicate that the transcriptional effects of aphidicolin are largely dependent on developmental context and do not support the existence of a conserved aphidicolin-induced transcriptional response in preimplantation embryos, consistent with previous analyses (Fig. S2I, J).

### Gene cluster functional enrichment reveals distinct responses to aphidicolin

To gain further insight into how blocking DNA replication during these early stages of mouse development alters transcription, we examined in greater detail the functional enrichments of genes grouped within the clusters described above (Tables S3, S4). As expected, enriched GO terms in the clusters that group genes unaffected by aphidicolin treatment (cluster #4 and #6), largely recapitulated those identified in the global analysis of up- or downregulated genes in each time window (Fig. S3A, B).

In the E2C-to-L2C time window, clusters #1 and #2, which contain genes whose expression is increased by aphidicolin treatment (Fig. 4E), shows enrichment for GO terms related to development and tissue regulation (Table S3). For example, cluster #1 includes genes encoding developmental transcription factors, such as *Foxd3*, *Gsc*, and *Tbx2*, and cluster #2 genes encoding components of signaling pathway, including *Dll4*, *Hes1* (Notch pathway) and *Ptch2* (Hedgehog pathway). On the other hand, in the 4C-to-8C window, cluster #1 was enriched for terms related to cell and tissue reorganization, whereas cluster #2 was enriched for various metabolic processes (Table S4). Nevertheless, cluster #1 also contained several genes encoding developmental regulators, similar to clusters #1 and #2 at the E2C-to-L2C time window. These included genes involved in early epiblast formation and gastrulation (*Eomes*, *Hesx1*, *Hhex*, *Nodal*, *Pou3f1*), or in the TE lineage (*Rhox6*, *Rhox9*, *Sfmbt2*). These findings suggest that blocking DNA replication drives premature activation of developmental and lineage-specifying genes during early cleavage stages.

Although clusters #3, #5 and # 7 from the E2C-to-L2C window contained too few genes for enrichment analysis (Table S1), manual inspection showed that genes from cluster #3 (genes whose increase from E2C-to-L2C is blocked by aphidicolin treatment; Fig. 4E) include several genes involved in cell cycle and mitosis (*Ccnf*, *Ckap2l*, *Kif11*) as well as other encoding chromosome-associated factors. This is also the case for cluster #5 (genes that do not change from E2C-to-L2C, but whose expression is downregulated by aphidicolin; *Cdca2*, *Ndc80*). In the 4C-to-8C time window, cluster #3 also includes developmental genes whose upregulation in blocked by aphidicolin (*Klf4*, *Klf5*, *Tbx3*, *Tcf3*, *Tfap2c*), and cluster #5 genes for DNA repair (*Recql*, *Top3a*, *Rcc1*) and chromatin regulation (*Suz12*, *Setd2*, *Kat5*, *Med1*).

Finally, cluster #8 from the E2C-to-L2C transition and #7 and #8 from the 4C-to-8C transition were enriched for GO categories related to metabolic processes (Tables S3, S4). These clusters contain genes that reduce their expression from E2C to L2C or from 4C to 8C, respectively, but whose downregulation is blocked by aphidicolin treatment (Fig. 4E). This observation led us to examine in more detail metabolic dysregulation occurring in preimplantation embryos in response to blocking DNA replication. KEGG pathway analysis by Gene Set Enrichment Analysis (GSEA) showed significant enrichments for metabolism associated categories in aphidicolin-treated embryos compared to age-matched 8C embryos (Fig. S4A). To further investigate this issue, we used METAFlux (Huang et al., 2023; Pan et al., 2025) to infer potential changes in metabolic fluxes from transcriptomic data based on gene expression levels (Fig. S5B, C; Table S5). Consistent with the transcriptomic analysis, aphidicolin-treated embryos exhibited relatively few metabolic alterations during the E2C-to-L2C transition (Fig. S4B). Notably, all predicted changes specifically induced by the presence of aphidicolin corresponded to increased fluxes (cluster vi, Fig. S5B), predominantly in lipid metabolic pathways, suggesting enhanced membrane lipid remodeling. This metabolic profile mirrors the transcriptional landscape and indicates that, although specific pathways may be affected by replication arrest, the acquisition of the L2C state is generally accompanied by a coordinated metabolic program that can be activated independently of DNA replication.

Interestingly, a markedly different scenario was observed during the 4C-to-8C transition. In contrast to the E2C-to-L2C window, numerous metabolic pathways that remained unchanged between control 4C and 8C embryos were significantly altered following aphidicolin treatment, with pathways being either downregulated or upregulated (clusters i′ and iv′, respectively; Fig. S5C). Specifically, pathways involved in energy production and cellular stress responses were upregulated, whereas anabolic processes associated with cell growth, proliferation, and differentiation appeared downregulated, as was also observed in the GO-term analysis of aphidicolin-treated embryos compared to 8C embryos (Fig. 4D, Table S2). These changes point to a substantial metabolic rewiring from a more proliferative developmental program toward a metabolically active state in response to replication stress. Together, these findings suggest that aphidicolin could elicit a robust stress response during the 4C-to-8C transition, that is not yet established at the 2-cell stage.

### Chromatin context drives susceptibility to inhibition of DNA replication

A possible explanation for the changes in gene expression observed upon aphidicolin treatment is that blocking DNA replication leads to alterations in chromatin organization in early embryos. To test this possibility, we examined the overlap and enrichment of different chromatin features among genes belonging to the expression clusters defined above (Fig. 4E).

Replication does not occur at random, but occurs in a defined temporal order that is cell-type specific, known as the replication timing (RT) program. This results in the partitioning of chromatin into early- and late-replicating domains, that are linked to three-dimensional genome organization and transcription. As such, early replicating domains are associated with active, transcribed chromatin, while late-replicating domains associate with silent heterochromatin (Vouzas and Gilbert, 2021). We took advantage of recent work that has mapped RT across mouse preimplantation development (Nakatani et al., 2024) to address the location of gene promoters from the previously defined clusters (Fig. 4E) to early- or late-replication domains. The majority of clusters in both time windows showed enrichment of genes in early-replicating domains (Fig. 5A), which is expected as transcriptomic analyses are inherently biased toward expressed genes. An exception to this pattern was cluster #1 from the E2C-to-L2C window, which is depleted in early-replicating domains throughout preimplantation development and in mouse embryonic stem cells (mESCs).

**Figure 5.**
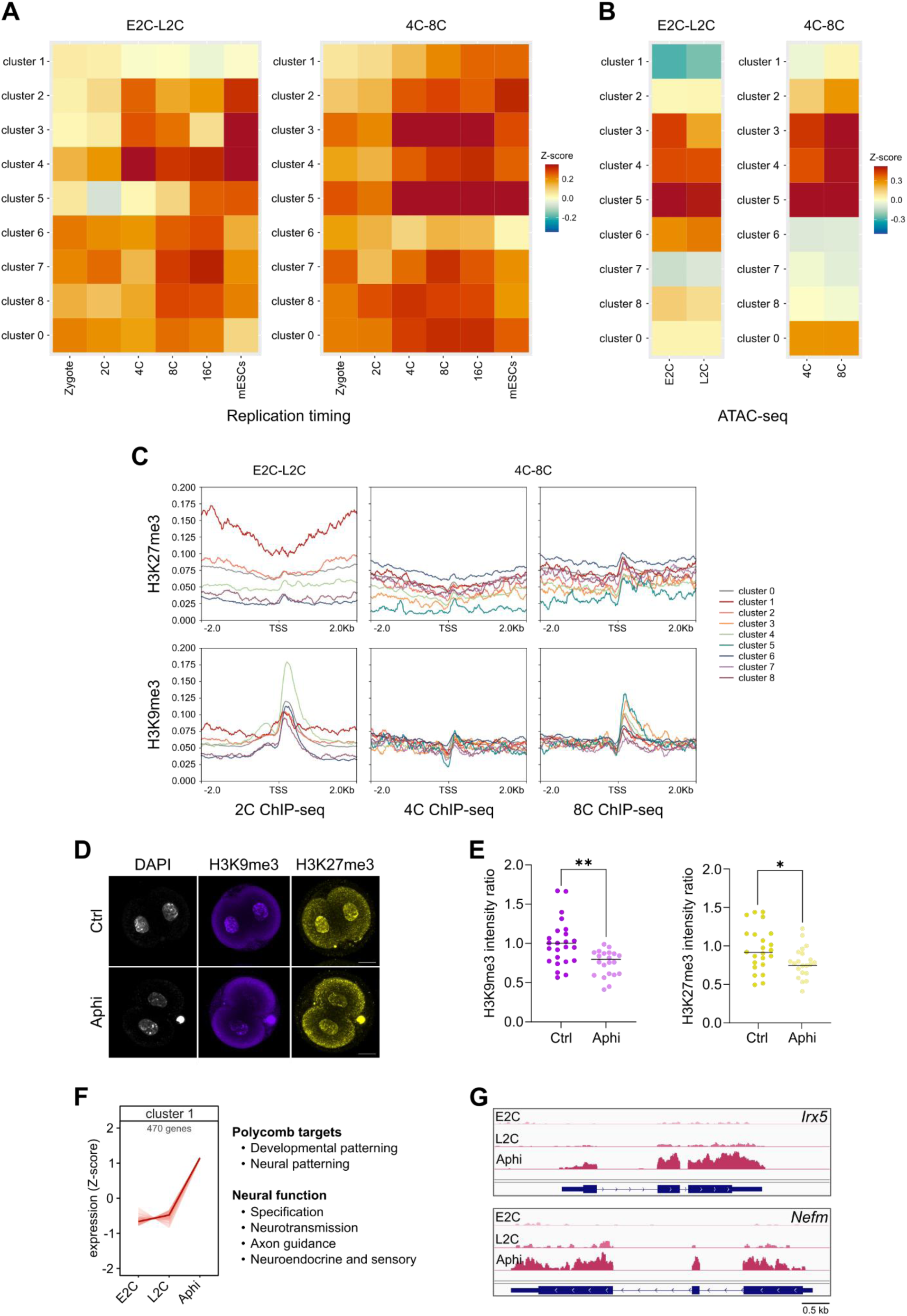
Blocking DNA replication alters the chromatin landscape. **(A)** Heatmaps showing the mean normalized enrichment (Z-score) of promoters of genes from each cluster (rows) from the E2C-to-L2C (left) and 4C-to-8C (right) time windows analyzed, in early replicating domains at the indicated developmental stages (columns). Color intensity indicates relative enrichment, with blue denoting lower and red higher enrichment. As the whole genome is divided into either early- or late-replicating domains, the values for late replicating domain enrichment are complementary to those of early replicating domains, and are therefore not shown. Replication timing data was obtained from Nakatani et al. (2024). **(B)** Heatmaps showing the mean normalized enrichment (Z-score) of promoters of genes from each cluster (rows) from the E2C-to-L2C (left) and 4C-to-8C (right) time windows analyzed, in previously described ATAC-seq peaks at the indicated developmental stages (columns). Color intensity indicates relative enrichment, with blue denoting lower and red higher enrichment. ATAC-seq data was obtained from Li et al. (2025). **(C)** Average profiles of H3K27me3 (top) and H3K9me3 (bottom) ChIP-seq signal on a 4 kilobase (kb) region surrounding the TSS of genes from each of the clusters defined in the E2C-to-L2C (left) and 4C-to-8C (middle and right) developmental windows (color code shown on the far right). Data for ChIP-seq generated in 2-cell (left), 4C (middle) and 8C (right) embryos for H3K27me3 was obtained from Liu et al. (2016) and for H3K9me3 from Wang et al. (2018). **(D)** Representative images of L2C control and age-matched aphidicolin-treated embryos labeled for total DNA (DAPI, gray, left column), H3K9me3 (blue, middle column) and H3K27me3 (yellow, right column). **(E)** Quantification of the fluorescent intensity as the ratio between the mean intensity of the nuclei from each embryo and the mean intensity of control embryos, in control L2C and age-matched aphidicolin-treated embryos, labeled with antibodies that recognize H3K9me3 (left, purple) or H3K27me3 (right, yellow). Each dot represents an individual nucleus (Ctrl, n=24; Aphi, n=21). * P<0.05; ** P<0.01; two-tailed Student’s t-test. **(F)** Expression trajectory of genes from cluster #1 of the E2C-to-L2C window (as in Fig. 4C). On the right, main categories of genes found in this cluster are indicated. **(G)** Representative examples of genes from cluster #1, showing RNA-seq reads mapped in the IGV viewer for the developmental patterning homeobox gene *Irx5* and the neurofilament medium chain coding gene *Nefm*.

To check chromatin accessibility of genes changing in our datasets, we used ATAC-seq information generated for the corresponding stages of mouse preimplantation development (Li et al., 2025). Overall, we observed a strong positive correlation between increasing expression and location of ATAC-seq peaks at gene transcriptional start sites (TSS), as a proxy for promoters (clusters #3 to #5 from both windows; Fig. 5B). In contrast, clusters that show a decrease in expression had less enrichment for ATAC-seq peaks (cluster #1, 4C-to-8C window; clusters #2, #6-#8 from both windows; Fig. 5B). Notably, cluster #1 from the E2C-to-L2C window stood out as being almost completely depleted from ATAC-seq peaks at promoters.

We next sought to analyze the epigenetic landscape associated to the different groups of the genes. To do so, we measured overlap of the TSS of genes from each cluster with different chromatin states defined in mESCs, based on pattens of histone modifications and chromatin-associated protein binding (Juan et al., 2016). The majority of clusters from both time windows showed enrichment for marks associated with active promoters and enhancers, states that were depleted of non-differential expressed genes (cluster #0) that showed marks for heterochromatin. Again, cluster #1 from the E2C-to-L2C window was the exception to this pattern: the TSSs of genes from this cluster were depleted from active promoter marks, instead showing high enrichment with marks associated with poised and repressed promoters (Fig. S5A). To obtain a more fine-grained view of these patterns, we analyzed enrichment of individual histone modifications and chromatin factors binding that underly the chromatin states described above (Fig. S5B) (Juan et al., 2016). Most interestingly, genes in cluster #1 from the E2C-to-L2C window were enriched in histone modifications (H2Aub1; H3K27me3) and chromatin factor binding (CBX7; EZH2, SUZ12, PHF19) characteristic of facultative heterochromatin mediated by Polycomb Repressive Complex 1 (PRC1) and Polycomb Repressive Complex 2 (PRC2), respectively. Furthermore, we also observed enrichment for H3K9me3, a marker of constitutive heterochromatin in cluster #1 genes.

### Regulation of genes located in broad Polycomb domains is disrupted by blocking DNA replication

The above observations strongly suggested that DNA replication was required to maintain, at least for a subset of loci, inactive heterochromatin domains during early mouse development. As our results suggest, aphidicolin treatment may disrupt this process, leading to the aberrant premature expression of genes that are normally silenced at these stages. To gain further insight into this process, we analyzed the distribution of histone modifications for both facultative PRC2-mediated heterochromatin (H3K27me3) and constitutive heterochromatin (H3K9me3) surrounding the promoter of genes from each cluster (Fig. 5C). For this, we took advantage of published datasets in mouse preimplantation development, using ChIP-seq data from 2-cell embryos for clusters from the E2C-to-L2C window, and data from 4C and 8C embryos for clusters from the 4C-to-8C window (Liu et al., 2016; Wang et al., 2018). As a control, we also examined the distribution of H3K4me3, that marks active promoters (Fig. S5C; Liu et al., 2016).

We observed that H3K27me3 was specifically enriched surrounding the TSSs of genes from cluster #1 of the E2C-to-L2C window, what is not the case for any other cluster of this window or for clusters from the 4C-to-8C window (Fig. 5C). This distribution differs from the canonical PRC2-mediated deposition of H3K27me3 at promoters, as can be seen emerging during the 4C-to-8C transition when 8C ChIP-seq data were used. However, it matches the non-canonical broad pattern of H3K27me3 distribution previously described in 2-cell preimplantation embryos (Zheng et al., 2016). On the other hand, H3K9me3 was enriched at gene promoters in both time windows (Fig. 5C). As expected for H3K4me3, a mark of active transcription, we observed the canonical enrichment at promoters, including the drop in signal at the nucleosome-depleted region at the TSS, with signal increasing during development (Fig. S5C).

The above observations suggested that inhibition of DNA replication may disrupt heterochromatin organization in the embryo. To test this, we quantified facultative and constitutive heterochromatin histone modifications (H3K27me3 and H3K9me3, respectively) by immunohistochemistry in L2C embryos and age-matched embryos treated with aphidicolin from the E2C stage (Fig. 5D). Indeed, we observed a consistent and significant reduction in both signals in aphidicolin-treated embryos (Fig. 5E), further supporting the functional link between replication and heterochromatin.

Taken together, this evidence points to genes we identified in cluster #1 from the E2C-to-L2C window (Fig. 5F) as showing a particular response to blocking of DNA replication. These genes are expressed at very low levels in E2C and L2C embryos, and aphidicolin treatment leads to their strong upregulation (Fig. 5G shows representative examples). Furthermore, they are preferentially located in late-replicating domains (Fig. 5A), not in regions of accessible chromatin (Figs 5B), and show chromatin features characteristic of Polycomb-regulated domains (Figs 5C, S5A, S5B). Initial functional annotation revealed that this cluster is enriched for developmental genes (Fig. 4C, Table S3), and a closer inspection showed that it included multiple patterning genes, particularly those involved in neural patterning. More surprisingly, it also included many genes involved in later neural function, such as neurotransmission, axon guidance, or neuroendocrine and sensory phenotypes (Fig. 5F, S5D). Consistent with these observations, many of these genes are known Polycomb-targets (Fig. S5D). Overall, these findings suggest that aphidicolin treatment disrupts the repression of developmentally regulated genes, particularly those subject to Polycomb-mediated silencing, this effect being significantly strong in the 2-cell stage.

### Developmental potential of preimplantation mouse embryos after replication arrest

Both changes to chromatin context and the observed consequences of transcriptional deregulation on metabolic pathways suggest that, although embryonic morphology and developmental progression appear largely preserved, the underlying viability of embryos may be perturbed. Furthermore, aphidicolin is known to induce replication stress and activate DNA damage response through ATM and ATR (Ozeri-Galai et al., 2008; Mazouzi et al., 2016). Despite not eliciting a common transcriptional response (Figs 4D, S2I, S2J), aphidicolin could still be having a deleterious effect on preimplantation embryos responsible for the main changes we observe. We first checked for specific transcriptional signatures of repair associated to DNA damage response (Wang and Wang, 2021) in our transcriptomic datasets, finding no significant changes induced by aphidicolin in E2C embryos (Table S1). As for embryos treated at 4C stage, we observed significant upregulation of *Cdkn1a*, *Mdmd2*, *Gadd45a*, *Bbc3*, and *Sesn2* (Table S1), what could be indicative of the activation of the ATM/p53 pathway. This is in line with the observation that double-stranded DNA damage response and the ATM pathway is not yet active in the very early mouse embryos, and only starts to function at 8C/morula stages (Yukawa et al., 2007; Wyatt et al., 2022).

To evaluate whether aphidicolin treatment and blocking DNA replication had long-term effects on embryonic viability and developmental potential, we treated E2C stage embryos with aphidicolin for 16 h, washed the drug, and allowed them to proceed for 24 or 48 h (Fig. 6A). After 24 h, control embryos had gone through two cleavage divisions, reaching the 8C stage, and begun to express differentiation markers such as GATA3 (although at low levels) and TEAD4 (Fig. 6B). Aphidicolin-treated embryos have a clear developmental delay, undergoing at most one division (Fig. 6C), and having approximately half the number of cells observed in control embryos (Fig. 6D). 48 h after drug removal, while controls expressed robustly GATA3 and TEAD4 in outer TE cells and had reached the late morula/early blastocyst stage (Fig. 6E-G), aphidicolin-treated embryos showed very poor survival (Fig. 6F, G). Nevertheless, the few embryos that progressed, despite being morphologically abnormal, initiated the expression of lineage differentiation markers (Fig. 6E).

**Figure 6.**
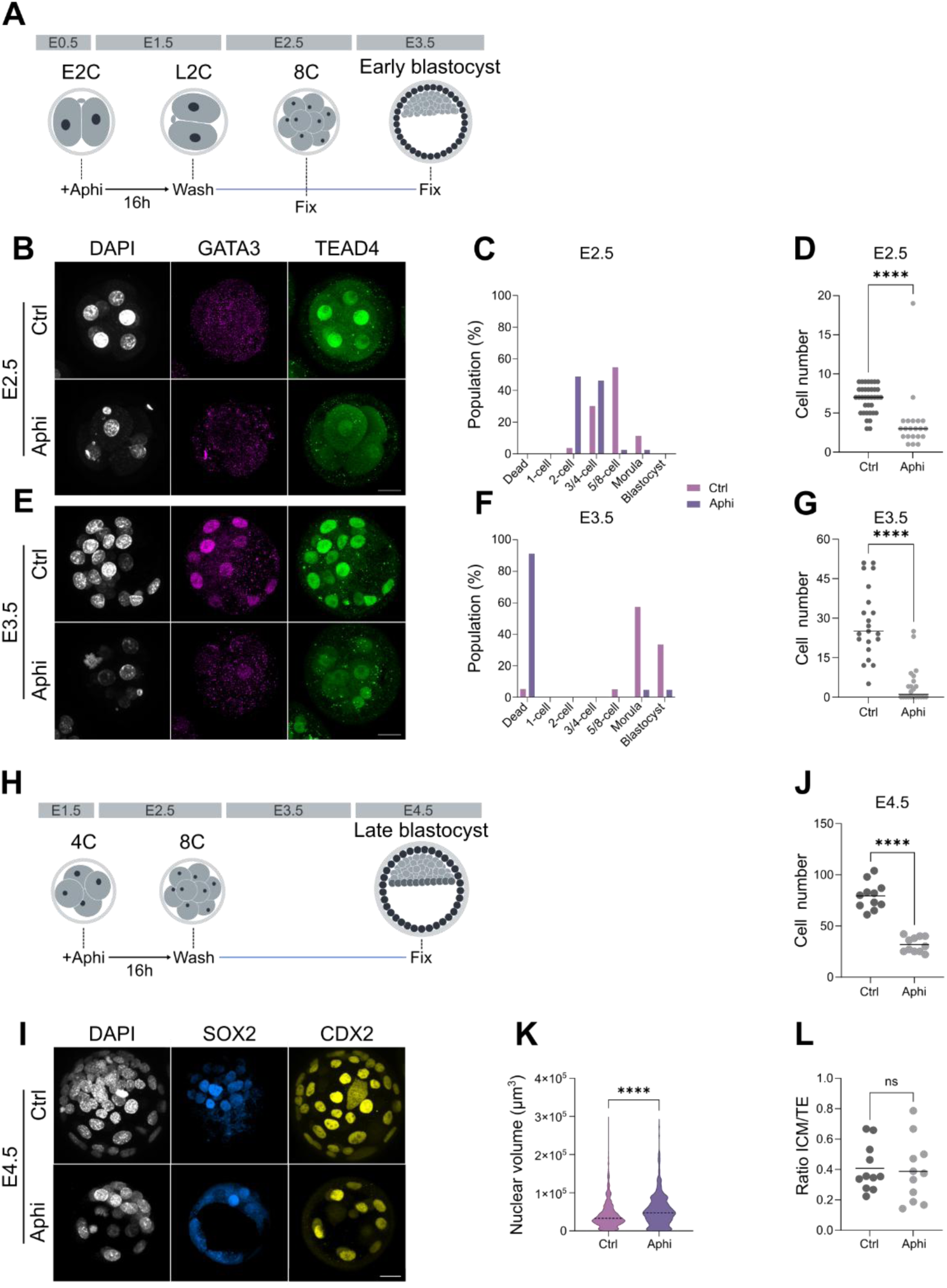
Recovery of embryonic development following aphidicolin treatment. **(A)** Experimental design showing the window of aphidicolin treatment from E2C to L2C stage, followed by washout and recovery for 24 or 48 h. Embryos were fixed when controls reached the 8C (24 h) or early blastocyst stage (48 h), respectively. **(B)** Representative images of control embryos (top) and embryos treated with aphidicolin from E2C to L2C stage, washed, and cultured for a further 48 h (bottom), corresponding to the time at which control embryos reach the 8C stage. Embryos are labeled with antibodies for GATA3 (middle, magenta) or TEAD4 (left, green). Nuclei are stained with DAPI (right, gray). **(C)** Proportion (as %) at each of the indicated developmental stages of control (n=53) and age-matched embryos treated with aphidicolin from E2C to L2C (n=41), washed, and left to develop until E2.5, when controls reached the 8C stage. **(D)** Number of cells at E2.5 of control (n=36) and age-matched embryos treated with aphidicolin from E2C to L2C (n=21). **** P<0.001; two-tailed Student’s t-test. **(E)** Representative images of control embryos (top) and embryos treated with aphidicolin from E2C to L2C stage, washed, and cultured for a further 48 h (bottom), corresponding to the time at which control embryos reached the early blastocyst stage. Embryos are labeled with antibodies for GATA3 (middle, magenta) or TEAD4 (left, green). Nuclei are stained with DAPI (right, gray). **(F)** Proportion (as %) at each of the indicated developmental stages of control (n=21) and age-matched embryos treated with aphidicolin from E2C to L2C (n=22), washed, and left to develop until E3.5, when controls reached the late morula – early blastocyst stage. **(G)** Number of cells at E3.5 of control (n=21) and age-matched embryos treated with aphidicolin from E2C to L2C (n=22). **** P<0.001; two-tailed Student’s t-test. **(H)** Experimental design showing the window of aphidicolin treatment from 4C to 8C stage, followed by washout and recovery for 48 h. Embryos were fixed when controls reached the late blastocyst stage. **(I)** Representative images of control embryos (top) and embryos treated with aphidicolin from 4C to 8C stage, washed, and cultured for a further 48 h (bottom), corresponding to the time at which control embryos reach the late blastocyst stage. Embryos are labeled with antibodies for SOX2 (middle, blue) or CDX2 (left, yellow). Nuclei are stained with DAPI (right, gray). **(J)** Number of cells at E4.5 of control (n=11) and age-matched embryos treated with aphidicolin from 4C to 8C (n=11). **** P<0.001; two-tailed Student’s t-test. **(K)** Quantification of the nuclear volume of cells from control (n=874 nuclei from 11 embryos) and age-matched embryos treated with aphidicolin from 4C to 8C (n=350 nuclei from 11 embryos), washed, and left to develop until E4.5, when controls reach the late blastocyst stage. **** P<0.001; two-tailed Student’s t-test. **(L)** Ratio of ICM to TE cells at E4.5, as identified by staining with SOX2 and CDX2 respectively, of control (n=11) and age-matched embryos treated with aphidicolin from 4C to 8C (n=11). ns, non-significant; two-tailed Student’s t-test.

However, when we treated embryos with aphidicolin at the 4C stage for 16 h, followed by drug washout and an additional 48 h of culture (Fig. 6H), viability was much higher. At this time point, control embryos had reached the late blastocyst stage, expressed distinct ICM and TE markers (SOX2 and CDX2, respectively; Fig. 6I) and contained approximately 80 cells (Fig. 6J). Following washout, aphidicolin-treated embryos developed into blastocysts expressing SOX2 and CDX2 (Fig. 6I), but of a smaller size and approximately half of the number of cells in controls (Fig. 6J). This is consistent with these embryos having skipped one cleavage division, but recovering developmental progression after that, in contrast to embryos treated at the 2-cell stage (Fig. 6A). We also observed an increase in nuclear volume in aphidicolin-treated embryos compared to age-matched controls (Fig. 6K), as we had when we analyzed embryos immediately after treatment (Fig. S1C). Finally, we wished to address if, despite their reduced cell number what could impact cell type proportions (Kukreja et al., 2024), aphidicolin-treated embryos maintained normal lineage allocation. To do so, we calculated the ratio of SOX2-positive cells to CDX2-positive cells in each embryo, and found no significant differences between treated embryos and controls (Fig. 6L). Thus, aphidicolin treatment at the 4C stage, inhibition of DNA replication, and the concomitant decrease in cell number did not significantly affect the relative proportions of the ICM and TE lineages.

## DISCUSSION

The first stages of mammalian development involve a small number of relatively slow cleavage divisions, from the one-cell embryo to the blastocyst, that occur alongside major transcriptional, epigenetic and morphological changes (Rossant, 2018; Burton and Torres-Padilla, 2025). Therefore, it has been difficult to determine the extent to which developmental progression depends on DNA replication and cell-cycle progression themselves. Here, we perturbed replication during the second and third embryonic cell cycles of the mouse and examined its consequences for initial transcription from the zygotic genome and for the first lineage decisions.

We observe that embryos in which DNA replication is blocked activate zygotic transcription, undergo compaction and polarization, initiate lineage specification programs and maintain proper gene expression trajectories despite failing to complete cell division. Nevertheless, DNA replication is required for the maintenance of specific repressive chromatin states, particularly at developmental loci associated with Polycomb-regulated heterochromatin (Zheng et al., 2016; Wang et al., 2018). Together, our results support a model in which developmental timing is largely uncoupled from cleavage progression, while replication contributes to the establishment of chromatin landscapes that restrict inappropriate developmental gene expression.

The observation that embryos continue to develop despite replication arrest builds on classical embryological studies, that showed how aphidicolin treatment does not block ZGA (Bolton et al., 1984; Aoki et al., 1997) or embryo progression (Smith and Johnson, 1985; Spindle et al., 1985). This suggested that early developmental events in mammals were governed by intrinsic programs rather than by cell number or mitotic history, what had also been observed in amphibians and insects (Newport and Kirschner, 1982; Edgar and O’Farrell, 1989), and more recently in nematodes or zebrafish (Nair et al., 2013; Kukreja et al., 2024). However, recent evidence from *Drosophila* shows that adult stem cells count the number of cell divisions to control multipotency (Tong et al., 2026). Here, we substantially extend these findings in the mouse by providing a genome-wide view of transcriptional progression during replication arrest, which show that aphidicolin-treated embryos more closely resemble age-matched controls than embryos with equivalent cell number. These results indicate that much of the transcriptional program operating during preimplantation development follows developmental time rather than replication or cell cycle progression, suggesting the existence of autonomous developmental timers that continue to operate in the absence of DNA synthesis.

A possible explanation could be that developmental timing at these stages is controlled by physical properties and morphological changes in the embryo, which may trigger signaling pathways involved in transcriptional responses and lineage decisions. This could be the case of the Hippo pathway, which has been involved in blastomere polarization and TE specification (Nishioka et al., 2009; Menchero et al., 2018). Consistent with this possibility, we observed that TEAD4, a key effector of Hippo signaling, is expressed at the right time in 4C embryos despite DNA replication block. Most intriguingly, the Hippo pathway has also been implicated in regulating ZGA (Yu et al., 2016; Zhu et al., 2020), a developmental transition during which we likewise observe transcriptional progression in the absence of DNA replication.

Despite the overall preservation of transcriptional programs during developmental progression, replication inhibition produced selective transcriptional changes that, importantly, did not cause a global collapse of transcription nor were characterized by a widespread aphidicolin-specific stress response. Instead, a relatively small subset of genes became aberrantly activated following replication arrest. These genes were highly enriched for later developmental regulators and patterning genes. So, our data suggest that the primary influence of replication on transcription during these stages may not be to promote activation, but rather to maintain repression of inappropriate transcriptional programs. Therefore, DNA replication would ensure that developmental regulators remain properly silenced until the appropriate embryonic stage is reached.

Several observations point toward chromatin organization as the underlying mechanism, mainly when we analyzed embryos from the E2C to L2C 2-cell stage. Genes most strongly affected by aphidicolin treatment were preferentially located in late-replicating domains (Nakatani et al., 2024), distant from accessible chromatin regions (Li et al., 2025) and enriched for chromatin signatures associated with Polycomb-mediated repression. This set of genes is normally transcriptionally repressed during ZGA and are enriched with broad H3K27me3 domains inherited from the oocyte or established de novo during cleavage stages and remain transcriptionally silent until post implantation development under normal conditions (Liu et al., 2016; Wang et al., 2018; Burton and Torres-Padilla, 2025; Matsuwaka et al., 2025). These genes include several *Hox* and *Irx* cluster genes, neurogenic fate determinants, and multiple signaling pathway genes involved in WNT, BMP, and TGF β signaling (i.e., *Wnt4*, *Dkk1*, *Bmp4*, or *Tgfb1*). Furthermore, inhibition of DNA replication reduced global H3K27me3 and H3K9me3 levels during the 2-cell stage. In this regard, it was recently shown that DNA replication was necessary for precise ZGA (Fan et al., 2026), arguing for subtle effects such as those we observe here that could be mediated by chromatin states. Also, studies in *C. elegans* showed that interfering with DNA replication during development led to de-repression of chromatin and anomalous gene expression (Klosin et al. 2017). Together, this suggest that replication contributes to the establishment or maintenance of heterochromatin during the extensive chromatin remodeling that accompanies ZGA. Failure to correctly shape these repressive domains may then permit premature transcription of developmental genes that should normally be maintained in a silent state.

Some questions remain unanswered. For example, why do we observe such a strong enrichment of neural-related genes? Importantly, this enrichment is not limited to neural developmental regulators and patterning genes, but also includes genes involved in establishing neuronal identity and terminal differentiation, such as those associated with synaptic transmission and axon guidance. These loci could be particularly dependent on replication-dependent chromatin assembly mechanisms. Alternatively, the activation of several neural developmental regulators in response to the block in DNA replication may be sufficient to drive the expression of downstream target genes involved in neuronal differentiation and cellular phenotypes. In any case, an intriguing observation is that this de-repression of late genes is observed only when DNA replication is blocked in the 2-cell embryos, but not at the 4C stage. This difference may relate to the progressive establishment of chromatin structure during early development (Du et al., 2022; Burton and Torres-Padilla, 2025), with earlier embryonic stages being less organized and therefore more susceptible to disruption upon replication block.

A second unexpected consequence of inhibiting DNA replication was the disruption of metabolic programs, particularly during the 4C to 8C transition. Computational analyses identified extensive alterations in pathways associated with lipid metabolism and, to a lesser extent, amino-acid metabolism. Whether these changes represent a direct consequence of replication arrest, a response to altered chromatin states, or an adaptive mechanism that supports embryo survival remains unclear. In this regard, it has been recently shown that lipid droplets have a fundamental role in early morphogenesis events in mouse preimplantation embryos (Aizawa et al., 2019; Mau et al., 2022). Therefore, these changes in lipid metabolism could be linked to ensuring correct development in spite of replication being inhibited.

Finally, the different outcomes we observe after release from aphidicolin treatment and replication arrest provide additional insight into stage-specific requirements for DNA replication. Embryos treated during the 2-cell stage showed poor long-term survival after drug washout, whereas embryos arrested during the 4C to 8C transition were capable of generating blastocysts with correct lineage determination and cell type proportions despite reduced cell numbers. This difference suggests that replication is particularly critical during the period surrounding ZGA, when chromatin architecture and epigenetic states are undergoing extensive reprogramming. By contrast, later stages appear more resilient to transient interruption of DNA synthesis once major aspects of genome organization have already been established. This would also be the case during zebrafish development, where DNA replication is dispensable or developmental progression at gastrulation (Kukreja et al., 2024). A possible explanation for our observations is the differential activity of DNA damage repair mechanisms during preimplantation development. It has been shown that ATM-dependent DNA repair is not active in the 2-cell embryo (Wyatt et al., 2022), therefore increasing lethality at this stage. On the other hand, ATR-mediated signaling induced by replication stress has been shown to promote totipotency in embryonic stem cells (Atashpaz et al., 2020). This could mean that aphidicolin-treated 2-cell embryos may be maintained in a totipotent or 2-cell-like state as a consequence of persistent replication stress, what would impair their ability to progress and therefore not survive after washout.

In summary, our results show that DNA replication is largely dispensable for early developmental and lineage-specification in the mouse embryo. Rather than acting as a primary driver of developmental progression, replication appears to allow specifically repressed chromatin states that preserve transcriptional fidelity during zygotic genome activation. These findings reveal the independence between developmental timing and cell-cycle progression, while highlighting DNA replication as a key contributor to the mechanisms that safeguard chromatin structure and correct developmental gene expression.

## MATERIALS AND METHODS

### Mouse embryo collection and culture

Female CD1 mice (6–8 weeks old) were superovulated by intraperitoneal injection of 5 IU equine chorionic gonadotropin (eCG; 0.15 mL) (LeonVet), followed 48 h later by 5 IU human chorionic gonadotropin (hCG) (Sigma, CG10). Females were immediately paired with CD1 males, and the presence of a vaginal plug the following morning was considered embryonic day 0.5 (E0.5). For zygote collection (E0.5), oviducts were dissected and the ampullae were mechanically opened with the aid of a 30G needle in M2 medium (Sigma, M7167) supplemented with hyaluronidase (Sigma, H3884) to release cumulus–oocyte complexes by a brief incubation. For collection of 4C stage embryos (E1.5), embryos were recovered by flushing M2 medium through the infundibulum of the oviduct using a filed 30G needle and a 1 ml syringe. Embryos were transferred with a fine glass pipette and cultured in KSOM medium (Sigma, MR-101-D) covered with mineral oil (Nanog Biotec, NO-400K) under standard conditions (37.5 °C and 5% CO2). Where indicated, embryos were cultured in KSOM supplemented with 5 µg/mL aphidicolin (Sigma, A0781) or DMSO (Sigma, D2650).

Mice were maintained at the CBM animal facility in Madrid, Spain, in accordance with national and European legislation. All animal procedures were approved by the CBM Animal Welfare Committee and the Regional Government of Madrid (ref. PROEX 116.3/25).

### EdU and EU staining

To assess genomic replication, E2C embryos were incubated with 20 μM 5-ethynyl-2’-deoxyuridine (EdU; 900584, Sigma-Aldrich) in the presence or absence of aphidicolin for 16 h. To examine active transcription during Zygotic Genome Activation (ZGA), zygotes and 2-cell embryos were incubated with 100 μM 5-ethynyl uridine (EU; Invitrogen, E10345) during the last 3 h of aphidicolin treatment. Following incubation, embryos were fixed in 4% paraformaldehyde (PFA; Electron Microscopy Sciences, 15710) for 10 min at room temperature (RT). Embryos were then briefly washed in 10% fetal bovine serum (FBS; Gibco, A5670701) diluted in PBS containing 0.1% Triton X-100 (0.1 % PBST; Sigma, X100), permeabilized with 0.5% PBST for 20 min at RT, and rinsed again in 10% FBS in 0.1% PBST. Samples were subsequently incubated in the Click-iT reaction cocktail, consisting of 80% PBST 0.1%, 10% 20 mM CuSO₄ (Sigma, C7631), 10% 0.5M L-ascorbic acid (Sigma, A4544), and 0.1% Alexa Fluor™ 647 azide (Thermofisher, A10277), for 30 min at RT. Finally, embryos were washed in 10% FBS in PBST 0.1% and incubated with DAPI for 10 min. Embryos were mounted in PBS in ibidi imaging chambers (Ibidi, 81507) and imaged using a Zeiss LSM 800 confocal microscope.

### Immunohistochemistry

Embryos at the desired stage were fixed in 4% PFA for 10 min at RT. Following fixation, embryos were permeabilized in 0.5% PBST for 15–30 min and blocked for 1 h at RT in 10% FBS diluted in 0.1 % PBST. Embryos were then incubated overnight at 4°C with the appropriate primary antibodies diluted in blocking solution. After washing in 0.1 % PBST, embryos were incubated with the corresponding secondary antibodies for 1 h at RT. Following additional washes in 0.1 % PBST, nuclei were counterstained with DAPI 5 µg/mL (Merck, 10236276001) for 10 min. The following antibodies and dilutions were used: rabbit polyclonal anti-GATA3 (H-48; Santa Cruz Biotechnology, sc-9009), 1:200; mouse monoclonal anti-TEAD4 (Abcam, ab58310), 1:200; rabbit polyclonal anti-pERM (Cell Signaling, 3141), 1:200; goat polyclonal anti-SOX2 (R&D Systems, AF2018), 1:100; mouse monoclonal anti-CDX2 (Abcam, ab86949), 2 drops; rabbit polyclonal anti-H3K9me3 (Abcam, ab8898), 1:200; mouse monoclonal anti-H3K27me3 (Abcam, ab6002), 1:200; Alexa Fluor 488 goat anti-Mouse IgG (Thermofisher, A-11001), 1:1000; Alexa Fluor 594 goat anti-Rabbit IgG (Thermofisher, A-11012), 1:1000; Alexa Fluor 568 donkey anti-Goat IgG (Thermofisher, A-11057), 1:1000; Alexa Fluor 647 chicken anti-Mouse IgG (Thermofisher, A-21463), 1:1000; Alexa Fluor 488 donkey anti-Rabbit IgG (Thermofisher, A-21206), 1:500; Alexa Fluor 647 donkey anti-Mouse IgG, (Thermofisher, A-31571), 1:500. Embryos were mounted in PBS in ibidi imaging chambers and imaged using a Zeiss LSM 800 confocal microscope.

### Image processing, quantification and statistical analysis

Confocal image processing was performed using Fiji (ImageJ; Schindelin et al., 2012; Schneider et al., 2012). Linear adjustments of brightness and contrast were applied equally to all images within the same experiment. Figures were assembled using Inkscape.

Image quantification was performed using one of two independent analysis pipelines, depending on the experiment. For experiments quantified using Arivis Vision4D (Zeiss), raw confocal images were first processed by applying background correction and denoising. Nuclei were subsequently automatically segmented based on the DAPI signal, and the mean fluorescence intensity of the indicated immunofluorescence channels was quantified within each segmented nucleus. The same analysis workflow and segmentation parameters were applied to all samples within each experiment to ensure consistency and reproducibility. Where indicated, fluorescence intensity values were normalized to the corresponding control condition and expressed as fluorescence intensity ratios, as specified in the relevant figure legends.

Alternatively, image analysis was performed using Fiji v1.54f (Schindelin et al., 2012; Schneider et al., 2012). Image quantification was carried out on raw images using custom Fiji macros to ensure identical analysis parameters across samples and reproducibility. Two-dimensional nuclear segmentation was performed using the StarDist deep-learning image segmentation plugin (Weigert et al., 2020). As the model had been trained on low-magnification images, immunofluorescence images were downscaled prior to segmentation, and the resulting regions of interest (ROIs) were subsequently rescaled to the original image dimensions for fluorescence intensity measurements. For three-dimensional analyses, DAPI image stacks were reordered to simulate 2D+T datasets, nuclei were segmented using the StarDist plugin and matched across adjacent optical sections using TrackMate. Volumetric cell measurements were obtained by combining nuclear segmentation with semi-automated whole-cell segmentation using the Segmentation Editor in Fiji. Quantification was performed using the 3D Manager implemented in the 3D ImageJ Suite (Ollion et al., 2013).

Graphical representation and statistical analyses were performed using GraphPad Prism (GraphPad Software, versions between 8.4.3 and 10.6.1). Details regarding sample size, biological replicates, definitions of individual data points, statistical analyses and significance used for each experiment are provided in the corresponding figure legends.

### RNAseq and computational analysis

RNA-seq was performed on four independent pools (replicates) of approximately 20–60 embryos each. RNA was extracted using the Arcturus PicoPure RNA Isolation Kit (Applied Biosystems, KIT0204). Library preparation and sequencing were carried out by the CRG Genomics Unit using a NextSeq 2000 platform, employing an ultra-low input protocol with poly(A) selection.

RNA-seq read sequences were assessed for quality using FastQC (www.bioinformatics.babraham.ac.uk/projects/fastqc) and mapped to the mm10 mouse reference genome with STAR (Dobin et al., 2013). BigWig tracks displayed in the IGV browser (Robinson et al. 2011) were computed with bamCoverage from deepTools (Ramirez et al., 2016) using -normalizeUsing CPM and -bs 1 options. Transcript abundance for protein-coding genes was quantified with featureCounts (Liao et al., 2014) based on the RefSeq annotation corresponding to the longest transcript isoform. Differential expression analysis across conditions was carried out using DESeq2 (Love et al., 2014). Genes with an adjusted p-value<0.01 and log_2_(fold-change)>|1| were classified as significantly differentially expressed genes (DEGs).

Genes with less than 30 total raw reads across the three conditions within each transition dataset were filtered out from further analyses. Hierarchical clustering of genes was performed based on their normalized Z-scores expression levels using the distance matrix and the hclust function in R. The number of clusters was determined to be 8 in both transition datasets using the clusGap function from cluster R package (Tibshirani et al., 2001). The Gap statistic method estimates the appropriate number of clusters by comparing the within-cluster variation of the observed data with that expected from randomly distributed data lacking cluster structure.

Gene Ontology (GO) enrichment analysis was conducted with the enrichGO function from the R package clusterProfiler (Yu et al., 2012) with default parameters and the “BP” (biological processes) ontology category. The most significant genes were ranked according to adjusted p-values for visualization. Overlap enrichment analysis of clusters across different chromatin signals was conducted using the crosswisePermTest function from the RegioneReloaded in R (Malinverni et al., 2023) with the following parameters: sampling=FALSE, genome=mm10, per.chromosome=TRUE, ranFUN=resampleRegions, evFUN=numOverlaps, ntimes=1000, universe - complete list of RefSeq genes. This function performs multiple permutation tests to assess the overlap between the gene list of interest and the genome coordinates of different chromatin signals, marks and states of chromatin previously defined. Histone metagene profiles were performed using computeMatrix and plotHeatmap functions from deepTools (Ramirez et al., 2016). Violin plots were generated with ggplot2 in R (Wickham 2016).

Metabolic fluxes were analyzed using METAFlux (Huang et al., 2023) based on the RPKM values of genes across conditions and a list of metabolites previously described in human blood and converted to mouse annotation using SBML2Flux (https://github.com/RestlessTail/SBML2Flux). Gene Set Enrichment Analysis (GSEA) (Subramanian et al., 2005) plots were perfomed based on the RPKM values of genes across replicates and using the KEGG database (Kanehisa and Goto, 2000).

### Analysis of public datasets

Promoter and gene body coordinates of genes within each cluster were intersected with previously described genomic and epigenetic features described in mouse embryos. These included open chromatin regions identified by scATAC-seq (Li et al. 2025), early replicating regions detected by scRepli-seq (Nakatani et al. 2024), and histone modifications mapped by ChIP-seq including H3K4me3 and H3K27me3 (Liu et al. 2016) and H3K9me3 (Wang et al. 2018). Additionally, chromatin states and histone mark annotations from mouse embryonic stem cells were obtained from Juan et al. (2016). Gene sets associated with minor and major ZGA, totipotency, and pluripotency used for expression analyses across developmental stages and experimental conditions were obtained from Yang et al. (2022). Gene sets affected by 24 h aphidicolin treatment in MEFs were obtained from Mazouzi et al. (2016). Published datasets were downloaded from GEO with accession numbers GSE45719 (scRNA-seq), GSE218365 (scRepli-seq), GSE73952 (H3K4me3 and H3K27me3 ChIP-seq), and GSE97778 (H3K9me3 ChIP-seq).

### Data and code availability

All scripts used are publicly available at https://github.com/aliciagallego/AphiProject and archived at Zenodo. RNA-seq data generated and analyzed in this work have been deposited in the NCBI Gene Expression Omnibus database (GEO; http://www.ncbi.nlm.nih.gov/geo/) under accession number GSE341830.

## ACKNOWLEDGEMENTS

We thank Maria Gomez (CBM), Barbara Pernaute (CABD), and present and past members of the Manzanares lab for constant encouragement and discussions; the CBM Advanced Light Microscopy Facility for excellent support with image acquisition and analysis; and the CBM Animal Facility for mice husbandry. This work was funded by grant PID2023-151742NB-I00 (MCIN/AEI/ 10.13039/501100011033. The Centro de Biología Molecular Severo Ochoa (CBM) is the recipient of a "Severo Ochoa Excellence” grant (CEX2021-001154-S) funded by MICIU/AEI (10.13039/501100011033) and receives institutional support from the Ramón Areces Foundation.

## AUTHOR CONTRIBUTIONS

Conceptualization: AAJ, AG, AB, MT, MM, MP; Data curation: AAJ, AG, MM, MP; Formal analysis: AAJ, AG. MP; Funding acquisition: MM; Investigation: AAJ, AG, EMB, AM, MP; Methodology: AAJ, AG, MP; Project administration: MM, MP; Software: AG, AB, MP: Supervision: MM, MP; Validation: AAJ, AG, MP; Visualization: AAJ, AG, MM, MP; Writing – original draft: MM, MP; Writing – review & editing: all authors.

## REFERENCES

Ahuja, A. K., Jodkowska, K., Teloni, F., Bizard, A. H., Zellweger, R., Herrador, R., Ortega, S., Hickson, I. D., Altmeyer, M., Mendez, J. and Lopes, M. (2016). A short G1 phase imposes constitutive replication stress and fork remodelling in mouse embryonic stem cells. Nat Commun 7, 10660.

Aizawa, R., Ibayashi, M., Tatsumi, T., Yamamoto, A., Kokubo, T., Miyasaka, N., Sato, K., Ikeda, S., Minami, N. and Tsukamoto, S. (2019). Synthesis and maintenance of lipid droplets are essential for mouse preimplantation embryonic development. Development 146, dev181925.

Al-Mousawi, J., Michetti, L., Castaldi, L., Liu, N., Villacorta, L., Benes, V. and Boskovic, A. (2026). High-resolution mapping of embryonic genome activation unveils a decoupling of transcription activation from precocious H3K4me3 removal. Sci Adv 12, eaec9545.

Aoki, F., Worrad, D. M. and Schultz, R. M. (1997). Regulation of transcriptional activity during the first and second cell cycles in the preimplantation mouse embryo. Dev Biol 181, 296–307.

Artus, J. and Cohen-Tannoudji, M. (2008). Cell cycle regulation during early mouse embryogenesis. Mol Cell Endocrinol 282, 78–86.

Atashpaz, S., Samadi Shams, S., Gonzalez, J. M., Sebestyen, E., Arghavanifard, N., Gnocchi, A., Albers, E., Minardi, S., Faga, G., Soffientini, E., Paolo Allievi, Cancila, V., Bachi, A., Fernández-Capetillo, Ó., Tripodo, C., Ferrari, F., López-Contreras, A. J. and Costanzo, V. (2020). ATR expands embryonic stem cell fate potential in response to replication stress. eLife 9, e54756.

Bolton, V. N., Oades, P. J. and Johnson, M. H. (1984). The relationship between cleavage, DNA replication, and gene expression in the mouse 2-cell embryo. Development 79, 139-163.

Burton, A. and Torres-Padilla, M. E. (2025). Epigenome dynamics in early mammalian embryogenesis. Nat Rev Genet 26, 587–603.

Chazaud, C. and Yamanaka, Y. (2016). Lineage specification in the mouse preimplantation embryo. Development 143, 1063–1074.

Ciemerych, M. A., Maro, B. and Kubiak, J. Z. (1999). Control of duration of the first two mitoses in a mouse embryo. Zygote 7, 293–300.

Collombet, S., Ranisavljevic, N., Nagano, T., Varnai, C., Shisode, T., Leung, W., Piolot, T., Galupa, R., Borensztein, M., Servant, N., Fraser, P., Ancelin, K. and Heard, E. (2020). Parental-to-embryo switch of chromosome organization in early embryogenesis. Nature 580, 142–146.

Dean, W. L. and Rossant, J. (1984). Effect of delaying DNA replication on blastocyst formation in the mouse. Differentiation 26, 134–137.

Dobin, A., Davis, C.A., Schlesinger, F., Drenkow, J., Zaleski, C., Jha, S., Batut, P., Chaisson, M. and Gingeras, T.R. (2013). STAR: ultrafast universal RNA-seq aligner. Bioinformatics 29, 15–21.

Du, Z., Zhang, K. and Wei Xie, W. (2022). Epigenetic reprogramming in early animal development. Cold Spring Harb Perspect Biol 14, a039677.

Eckersley-Maslin, M. A., Alda-Catalinas, C. and Reik, W. (2018). Dynamics of the epigenetic landscape during the maternal-to-zygotic transition. Nat Rev Mol Cell Biol 19, 436–450.

Edgar, B. A. and O’Farrell, P. H. (1989). Genetic control of cell division patterns in the Drosophila embryo. Cell 57, 177–187.

Fan, Q., Wu, X., Dang, Y., Dong, L., Wang, W., Kong, F., Wang, L., Lu, X., Liu, B., Ji, S. and Xie, X. (2026). THAP1 is a maternal effect factor required for the first cell cycle via Rrm1 in early mouse embryos. EMBO Rep 27, 1813–1829.

Home, P., Ray, S., Dutta, D., Bronshteyn, I., Larson, M. and Paul, S. (2009). GATA3 is selectively expressed in the trophectoderm of peri-implantation embryo and directly regulates Cdx2 gene expression. J Biol Chem 284, 28729–28737.

Huang, Y., Mohanty, V., Dede, M., Tsai, K., Daher, M., Li, L., Rezvani, K. and Chen, K. (2023). Characterizing cancer metabolism from bulk and single-cell RNA-seq data using METAFlux. Nat Commun 14, 4883.

Ikegami, S., Taguchi, T., Ohashi, M., Oguro, M., Nagano, H. and Mano, Y. (1978). Aphidicolin prevents mitotic cell division by interfering with the activity of DNA polymerase-α. Nature 275, 458–460.

Juan, D., Perner, J., Carrillo de Santa Pau, E., Marsili, S., Ochoa, D., Chung, HR., Vingron, M., Rico, D. and Valencia, A. (2016). Epigenomic co-localization and co-evolution reveal a key role for 5hmC as a communication hub in the chromatin network of ESCs. Cell Rep 14, 1246–1257.

Kanehisa, M. and Goto, S. (2000). KEGG: kyoto encyclopedia of genes and genomes. Nucl Acids Res 28, 27–30.

Ke, Y., Xu, Y., Chen, X., Feng, S., Liu, Z., Sun, Y., Yao, X., Li, F., Zhu, W., Gao, L., Chen, H., Du, Z., Xie, W., Xu, X., Huang, X. and Liu, J. (2017). 3D chromatin structures of mature gametes and structural reprogramming during mammalian embryogenesis. Cell 170, 367–381.

Klosin, A., Reis, K., Hidalgo-Carcedo, C., Casas, E., Vavouri, T. and Lehner, B. (2017). Impaired DNA replication derepresses chromatin and generates a transgenerationally inherited epigenetic memory. Sci Adv 3, e1701143.

Kukreja, K., Jia, B. Z., McGeary, S. E., Patel, N., Megason, S. G. and Klein, A. M. (2024). Cell state transitions are decoupled from cell division during early embryo development. Nat Cell Biol 26, 2035–2045.

Li, M., Jiang, Z., Xu, X., Wu, X., Liu, Y., Chen, K., Liao, Y., Li, W., Wang, X., Guo, Y., Zhang, B., Wen, L., Kee, K. and Tang, F. (2025). Chromatin accessibility landscape of mouse early embryos revealed by single-cell NanoATAC-seq2. Science 387, eadp4319.

Liao, Y., Smyth, G. K. and Shi, W. (2014). featureCounts: an efficient general-purpose program for assigning sequence reads to genomic features. Bioinformatics 30, 923–930.

Liu, X., Wang, C., Liu, W., Li, J., Li, C., Kou, X., Chen, J., Zhao, Y., Gao, H., Wang, H., Zhang, Y., Gao, Y., Gao, S. (2016). Distinct features of H3K4me3 and H3K27me3 chromatin domains in pre-implantation embryos. Nature 537, 558–562.

Louvet, S., Aghion, J., Santa-Maria, A., Mangeat, P. and Maro, B. (1996). Ezrin becomes restricted to outer cells following asymmetrical division in the preimplantation mouse embryo. Dev Biol 177, 568–579.

Love, M. I., Huber, W. and Anders, S. (2014). Moderated estimation of fold change and dispersion for RNA-seq data with DESeq2. Genome Biol 15, 550.

Malinverni, R., Corujo, D., Gel, B. and Buschbeck, M. (2023). regioneReloaded: evaluating the association of multiple genomic region sets. Bioinformatics 39, btad704.

Matsuwaka, M., Kumon, M. & Inoue, A. (2025). H3K27 dimethylation dynamics reveal stepwise establishment of facultative heterochromatin in early mouse embryos. Nat Cell Biol 27, 28–38.

Mau, K. H. T., Karimlou, D., Barneda, D., Brochard, V., Royer, C., Leeke, B., de Souza, R. A., Pailles, M., Percharde, M., Srinivas, S., Jouneau, A., Christian, M. and Azuara, V. (2022). Dynamic enlargement and mobilization of lipid droplets in pluripotent cells coordinate morphogenesis during mouse peri-implantation development. Nat Commun 13, 3861.

Mazouzi, A., Stukalov, A., Müller, A. C., Chen, D., Wiedner, M., Prochazkova, J., Chiang, S. C., Schuster, M., Breitwieser, F. P., Pichlmair, A., El-Khamisy, S. F., Bock, C., Kralovics, R., Colinge, J., Bennett, K. L. and Loizou, J. I. (2016). A comprehensive analysis of the dynamic response to aphidicolin-mediated replication stress uncovers targets for ATM and ATMIN. Cell Rep 15, 893–908.

Menchero, S., Sainz de Aja, J. and Manzanares, M. (2018). Our first choice: cellular and genetic underpinnings of trophectoderm identity and differentiation in the mammalian embryo. Curr Top Dev Biol 128, 59–80.

Nair, G., Walton, T., Murray, J. I. and Raj, A. (2013). Gene transcription is coordinated with, but not dependent on, cell divisions during C. elegans embryonic fate specification. Development 140, 3385–3394.

Nakatani, T., Lin, J., Ji, F., Ettinger, A., Pontabry, J., Tokoro, M., Altamirano-Pacheco, L., Fiorentino, J., Mahammadov, E., Hatano, Y., Van-Rechem, C., Chakraborty, D., Ruiz-Morales, E. R., Arguello Pascualli, P. Y., Scialdone, A., Yamagata, K., Whetstine, J. R., Sadreyev, R. I. and Torres-Padilla, M.-E. (2022). DNA replication fork speed underlies cell fate changes and promotes reprogramming. Nat Genet 54, 318–327.

Nakatani, T., Schauer, T., Altamirano-Pacheco, L., Klein, K. N., Ettinger, A., Pal, M., Gilbert, D. M. and Torres-Padilla, M.-E. (2024). Emergence of replication timing during early mammalian development. Nature 625, 401–409.

Newport, J. and Kirschner, M. (1982). A major developmental transition in early Xenopus embryos: I. characterization and timing of cellular changes at the midblastula stage. Cell 30, 675–686.

Nishioka, N., Yamamoto, S., Kiyonari, H., Sato, H., Sawada, A., Ota, M., Nakao, K. and Sasaki, H. (2008). Tead4 is required for specification of trophectoderm in pre-implantation mouse embryos. Mech Dev 125, 270–283.

Nishioka, N., Inoue, K., Adachi, K., Kiyonari, H., Ota, M., Ralston, A., Yabuta, N., Hirahara, S., Stephenson, R. O., Ogonuki, N., Makita, R., Kurihara, H., Morin-Kensicki, E. M., Nojima, H., Rossant, J., Nakao, K., Niwa, H. and Sasaki, H. (2009). The Hippo signaling pathway components Lats and Yap pattern Tead4 activity to distinguish mouse trophectoderm from inner cell mass. Dev Cell 16, 398–410.

Ollion, J., Cochennec, J., Loll, F., Escudé, C. and Boudier, T. (2013). TANGO: a generic tool for high-throughput 3D image analysis for studying nuclear organization. Bioinformatics 29, 1840–1841.

Ozeri-Galai E, Schwartz M, Rahat A, Kerem B. (2008). Interplay between ATM and ATR in the regulation of common fragile site stability. Oncogene 27, 2109–2117.

Pan, Y., Huang, Y., Mohanty, V. and Chen, K. (2025). Inferring Metabolic Flux from Gene Expression Data Using METAFlux. Methods Mol Biol 2932, 187–202.

Pratt, H. P, Chakraborty, J. and Surani, M. A. (1981) Molecular and morphological differentiation of the mouse blastocyst after manipulations of compaction with cytochalasin D. Cell 26, 279–292.

Ramírez, F., Ryan, D.P., Grüning, B., Bhardwaj, V., Kilpert, F., Richter, A.S., Heyne, S., Dündar, F., Manke, T. (2016). deepTools2: a next generation web server for deep-sequencing data analysis. Nucl Acids Res 44, W160–W165.

Robinson, J.T., Thorvaldsdóttir, H., Winckler, W., Guttman, M., Lander, E.S., Getz, G. and Mesirov, J.P. (2011). Integrative genomics viewer. Nat Biotechnol 29, 24–26.

Rollins, M. B. and Andrews, M. T. (1991). Morphogenesis and regulated gene activity are independent of DNA replication in Xenopus embryos. Development 112, 559–569.

Rossant, J. (2018). Genetic Control of Early Cell Lineages in the Mammalian Embryo. Annu Rev Genet 52, 185–201.

Schindelin, J., Arganda-Carreras, I., Frise, E., Kaynig, V., Longair, M., Pietzsch, T., Preibisch, S., Rueden, C., Saalfeld, S., Schmid, B., Tinevez, J. Y., White, D. J., Hartenstein, V., Eliceiri, K., Tomancak, P. and Cardona, A. (2012). Fiji: an open-source platform for biological-image analysis. Nat Methods 9, 676–682.

Schneider, C. A., Rasband, W. S. and Eliceiri, K. W. (2012). NIH Image to ImageJ: 25 years of image analysis. Nat Methods 9, 671–675.

Smith, R. K. and Johnson, M. H. (1985). DNA replication and compaction in the cleaving embryo of the mouse. J Embryol Exp Morphol 89, 133–148.

Spadari, S., Sala, F. and Pedrali-Noy, G. (1982). Aphidicolin: A specific inhibitor of nuclear DNA replication in eukaryotes. Trends Biochem Sci 7, 29–32.

Spindle, A., Nagano, H. and Pedersen, R. A. (1985). Inhibition of DNA replication in preimplantation mouse embryos by aphidicolin. J Exp Zool 235, 289–295.

Subramanian, A., Tamayo, P., Mootha, V. K., Mukherjee, S., Ebert, B. L., Gillette, M. A., Paulovich, A., Pomeroy, S. L, Golub, T. R., Lander, E. S. and Mesirov, J. P. (2005). Gene set enrichment analysis: A knowledge-based approach for interpreting genome-wide expression profiles. Proc Natl.Acad.Sci.USA.102, 15545-15550.

Tibshirani, R., Walther, G. and Hastie, T. (2001). Estimating the number of clusters in a data set via the gap statistic. J R Stat Soc Series B Stat Methodol 63, 411–423.

Tong, D., Li, A., Jiang, Q., Yuan, Q., Liu, X., He, X., Liang, J., Han, Y. and Guo, Z. (2026). Intestinal stem cells count self-renewal divisions to switch multipotency. Nature, doi: 10.1038/s41586-026-10814-y.

Vouzas, A. E. and Gilbert, D. M. (2021). Mammalian DNA replication timing. Cold Spring Harb Perspect Biol 13, a040162.

Wang, C., Liu, X., Gao, Y., Yang, L., Li, C., Liu, W., Chen, C., Kou, X., Zhao, Y., Chen, J., Wang, Y., Le, R., Wang, H., Duan, T., Zhang, Y. and Gao, S. (2018). Reprogramming of H3K9me3-dependent heterochromatin during mammalian embryo development. Nat Cell Biol 20, 620–631.

Wang, X. and Wang, S.M. (2021). DNA damage repair system in C57BL/6 J mice is evolutionarily stable. BMC Genomics 22, 669.

Weigert, M., Schmidt, U., Haase, R., Sugawara, K. and Myers, G. (2020). Star-convex polyhedra for 3D object detection and segmentation in microscopy. The IEEE Winter Conference on Applications of Computer Vision (2020), 3666-3673.

Wickham H (2016). ggplot2: elegant graphics for data analysis. Springer-Verlag New York. ISBN 978–3-319-24277-4.

Wyatt, C. D., Pernaute, B., Gohr, A., Miret-Cuesta, M., Goyeneche, L., Rovira, Q., Salzer, M. C., Boke, E., Bogdanovic, O., Bonnal, S. and Irimia, M. (2022). A developmentally programmed splicing failure contributes to DNA damage response attenuation during mammalian zygotic genome activation. Sci Adv 8, eabn4935.

Xu, S., Wang, N., Zuccaro, M. V., Gerhardt, J., Iyyappan, R., Scatolin, G. N., Jiang, Z., Baslan, T., Koren, A. and Egli, D. (2024). DNA replication in early mammalian embryos is patterned, predisposing lamina-associated regions to fragility. Nat Commun 15, 5247.

Yang, M., Yu, H., Yu, X., Liang, S., Hu, Y., Luo, Y., Izsvák, Z., Sun, C. and Wang, J. (2022). Chemical-induced chromatin remodeling reprograms mouse ESCs to totipotent-like stem cells. Cell Stem Cell 29, 400–418.

Yu, G., Wang, L. G., Han, Y. and He, Q. Y. (2012). clusterProfiler: an R package for comparing biological themes among gene clusters. OMICS 16, 284–287.

Yu, C., Ji, S.-Y., Dang, Y.-J., Sha, Q.-Q., Yuan, Y.-F., Zhou, J.-J., Yan, L.-Y., Qiao, J., Tang, F. and Fan, H.-Y. (2016). Oocyte-expressed yes-associated protein is a key activator of the early zygotic genome in mouse. Cell Res 26, 275–287.

Yukawa, M., Oda, S., Mitani, H., Nagata, M. and Aoki, F. (2007). Deficiency in the response to DNA double-strand breaks in mouse early preimplantation embryos. Biochem Biophys Res Commun 358, 578–584.

Zeman, M. K. and Cimprich, K. A. (2014). Causes and consequences of replication stress. Nat Cell Biol 16, 2–9.

Ziomek, C. and Johnson, M. (1980). Cell surface interaction induces polarization of mouse 8C blastomeres at compaction. Cell 21, 935–942.

Zheng, H., Huang, B., Zhang, B., Xiang, Y., Du, Z., Xu, Q., Li, Y., Wang, Q., Ma, J., Peng, X., Xu, F. and Xie, W. (2016). Resetting epigenetic memory by reprogramming of histone modifications in mammals. Mol Cell 63, 1066–1079.

Zhu, M., Cornwall-Scoones, J., Wang, P., Handford, C. E., Na, J., Thomson, M. and Zernicka-Goetz, M. (2020). Developmental clock and mechanism of de novo polarization of the mouse embryo. Science 370, eabd2703.

