## Supplemental material for "DNA replication is dispensable for developmental progression, but required for heterochromatin organization at mouse zygotic genome activation"

**Arroyo-Jimenez et al.**

### Supplemental Figures S1-S5

### Supplemental Tables S1-S5

**Table S1.** Differential gene expression analysis in the E2C-L2C and in the 4C-8C time windows.

**Table S2.** Gene Ontology (GO) enrichment analysis (Biological Processes) of differentially expressed genes in the E2C-L2C and in the 4C-8C time windows.

**Table S3.** Gene Ontology (GO) enrichment analysis (Biological Processes) of gene clusters from the E2C-L2C time window.

**Table S4.** Gene Ontology (GO) enrichment analysis (Biological Processes) of gene clusters from the 4C-8C time window.

**Table S5.** METAFlux analysis of transcriptional changes in the E2C-L2C and in the 4C-8C time windows.

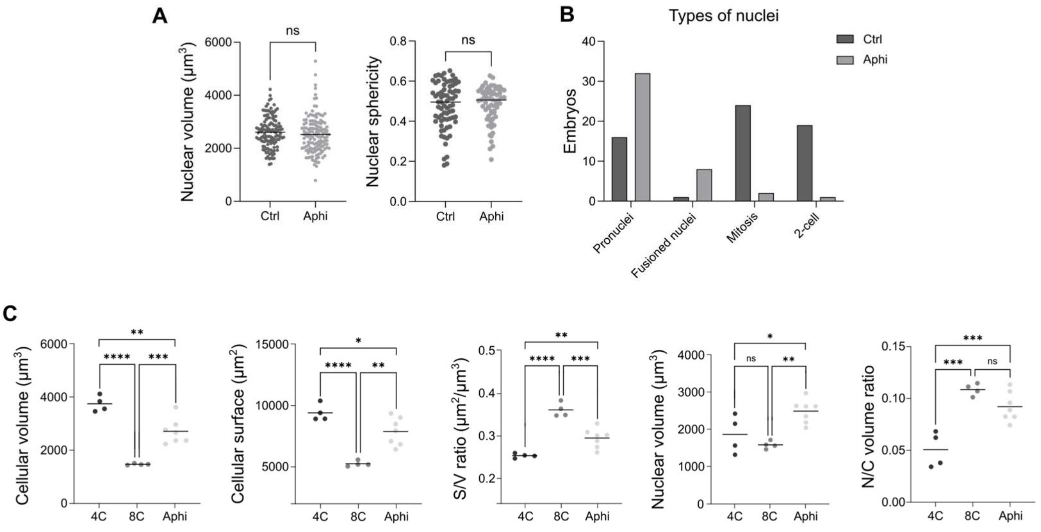


**Figure S1. Developmental effects of the inhibition of DNA replication in preimplantation embryos.**

**(A)** Quantification of nuclear volume (left) and nuclear sphericity (right) in control L2C embryos and age-matched aphidicolin-treated embryos from the E2C stage. Each dot represents an individual embryo (n=138, 228 nuclei quantified), and horizontal lines indicate the mean. * P<0.05; ns, non-significant; two-tailed Student's t-test.

**(B)** Distribution of control E2C (n=36) and age-matched embryos treated with aphidicolin from one-cell to E2C (n=41), according to developmental progression: embryos with two visible pronuclei, fused pronuclei, a nucleus undergoing mitosis, or two-cell embryos (left to right).

**(C)** Morphometric measurements of control 4-cell and 8-cell embryos, and age-matched aphidicolin-treated embryos. Each dot represents an individual embryo (n=4, 4-cell; n=4, 8-cell; n=7, Aphi). * P<0.05; *** P<0.005; **** P<0.001; ns, non-significant; two-tailed Student's t-test.

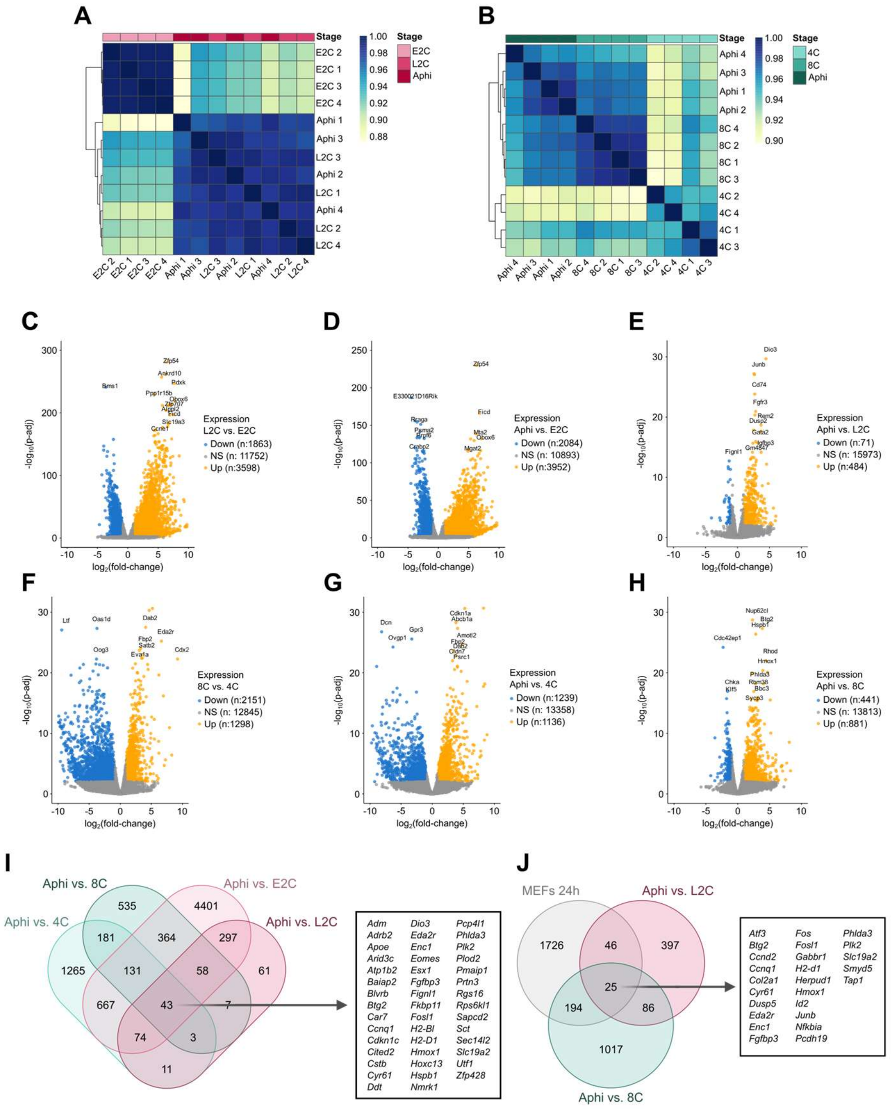


**Figure S2. Transcriptional changes in response to inhibition of DNA replication in preimplantation embryos.**

**(A)** Hierarchical clustering heatmap of the E2C-to-L2C RNA-seq dataset, showing the similarity among samples based on pairwise correlation values. Darker blue colors indicate higher correlation coefficients, whereas lighter shades represent lower correlations. Samples cluster into two major groups: the E2C samples (E2C1–E2C4) and the Aphi/L2C samples (Aphi1–Aphi4 and L2C1–L2C4), demonstrating that global transcription is largely maintained after DNA replication arrest.

**(B)** Hierarchical clustering heatmap of the 4C-to-8C RNA-seq dataset, based on pairwise correlation values. Darker blue colors indicate higher correlation coefficients, whereas lighter shades represent lower correlations. Samples cluster into two major groups: the 4C samples (4C1–4C4) and the Aphi/8C samples (Aphi1–Aphi4 and 8C1–8C4), demonstrating that global transcription is largely maintained after DNA replication arrest.

**(C-H)** Volcano plots showing differential gene expression between groups from the E2C-to-L2C RNA-seq dataset (**C**-**E**), and the 4C-to-8C dataset (**F**-**H**). Each point represents a gene, plotted according to its log2 fold change on the x-axis and -log10 adjusted P-value on the y-axis. Significantly upregulated genes are shown in orange, downregulated genes in blue, and non-differentially expressed genes in gray. Selected differentially expressed genes for each comparison are labeled, and number of genes in each category are indicated.

**(I)** Venn diagram showing the overlap of DEGs between embryos treated with aphidicolin at E2C stage for 16 h compared with either E2C (aphi-E2C) or L2C (aphi-L2C) control embryos, and embryos treated with aphidicolin at 4-cell stage for 16 h compared with either 4-cell (aphi-4C) or 8-cell (aphi-8C) control embryos. Genes common to all four sets (n=45) are listed on the left.

**(J)** Venn diagram showing the overlap of DEGs in MEFs treated with aphidicolin for 24 h (mefs 24h) and DEGs between embryos treated with aphidicolin at E2C stage for 16 h compared with L2C control embryos (aphi-L2C), and embryos treated with aphidicolin at 4-cell stage for 16 h compared with 8-cell control embryos (aphi-8C). Genes common to all three sets (n=25) are listed on the left. Gene expression data of MEFs was obtained from Mazouzi et al. (2016).

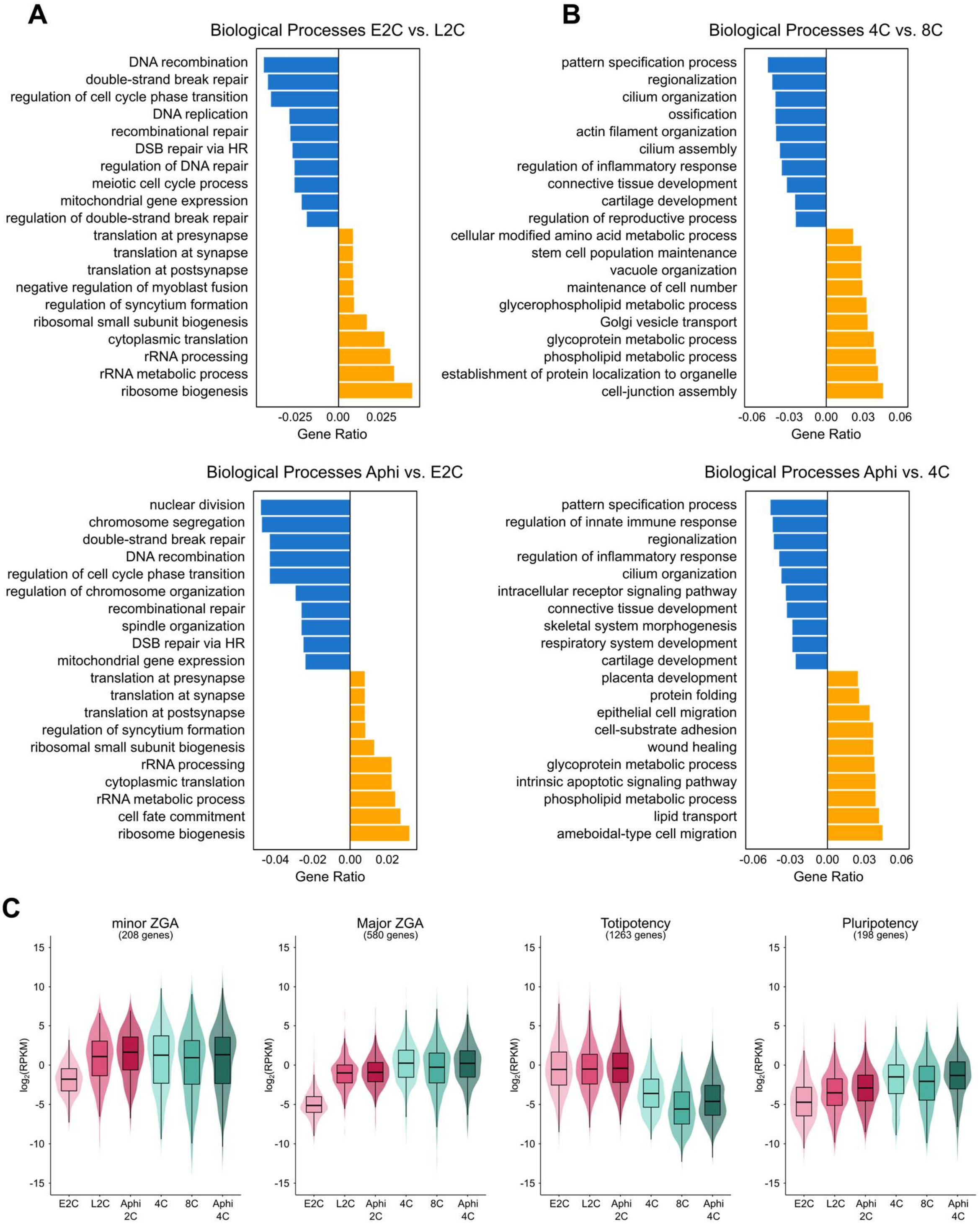


**Figure S3. Functional enrichment of differentially expressed genes.**

**(A)** GO Biological Process enrichment analysis of up (orange) and down (blue) DEGs between L2C and E2C embryos (top) and between aphidicolin-treated embryos at the E2C stage for 16 h and E2C control embryos (bottom). Top ten statistically significant categories (adj p-value<0,01) are shown, ranked by gene ratio.

**(B)** GO Biological Process enrichment analysis of up (orange) and down (blue) DEGs between 8C and 4C embryos (top) and between aphidicolin-treated embryos at the 4C stage for 16 h and 8C control embryos (bottom). Top ten statistically significant categories (adj p-value<0,01) are shown, ranked by gene ratio.

**(C)** Violin plots of log2 (RPKM) expression values for minor ZGA, major ZGA, totipotency, and pluripotency gene sets (Yang et al., 2022) in E2C, L2C, Aphi E2C-L2C, 4-cell, 8-cell, and Aphi 4C-8C RNA-seq samples. The width of each violin indicates the density of genes at a given expression level, horizontal lines the median expression value, and boxplots 25th to 75th percentiles of expression.

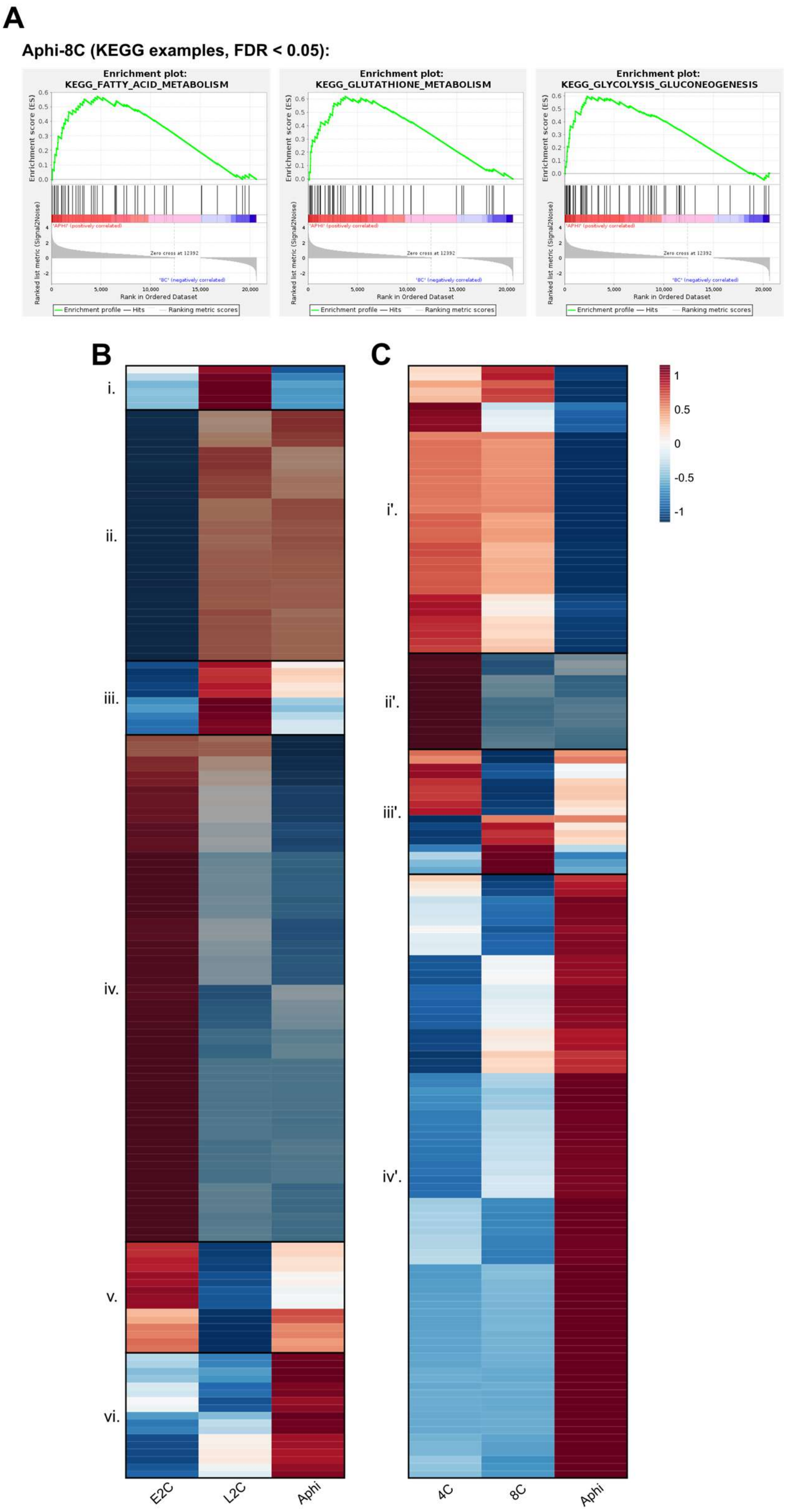


**Figure S4. Metabolic dysregulation in response to inhibition of DNA replication.**

**(A)** Representative GSEA plots for significantly enriched KEGG pathways (FDR < 0.05) in the 4C-to-8C dataset, including fatty acid metabolism, glutathione metabolism, and glycolysis/gluconeogenesis.

**(B-C)** METAFlux spearman correlation heatmaps of metabolic fluxes across samples in the E2C-to-L2C (**A**) and the 4C-to-8C (**B**) RNA-seq datasets. Rows represent fluxes, and columns experimental groups. Correlation values are displayed using a blue-red color scale, where red indicates higher and blue indicates lower correlation. Distinct clusters of features with similar expression patterns are separated by black borders. Clusters that do not change upon aphidicolin treatment are shaded.

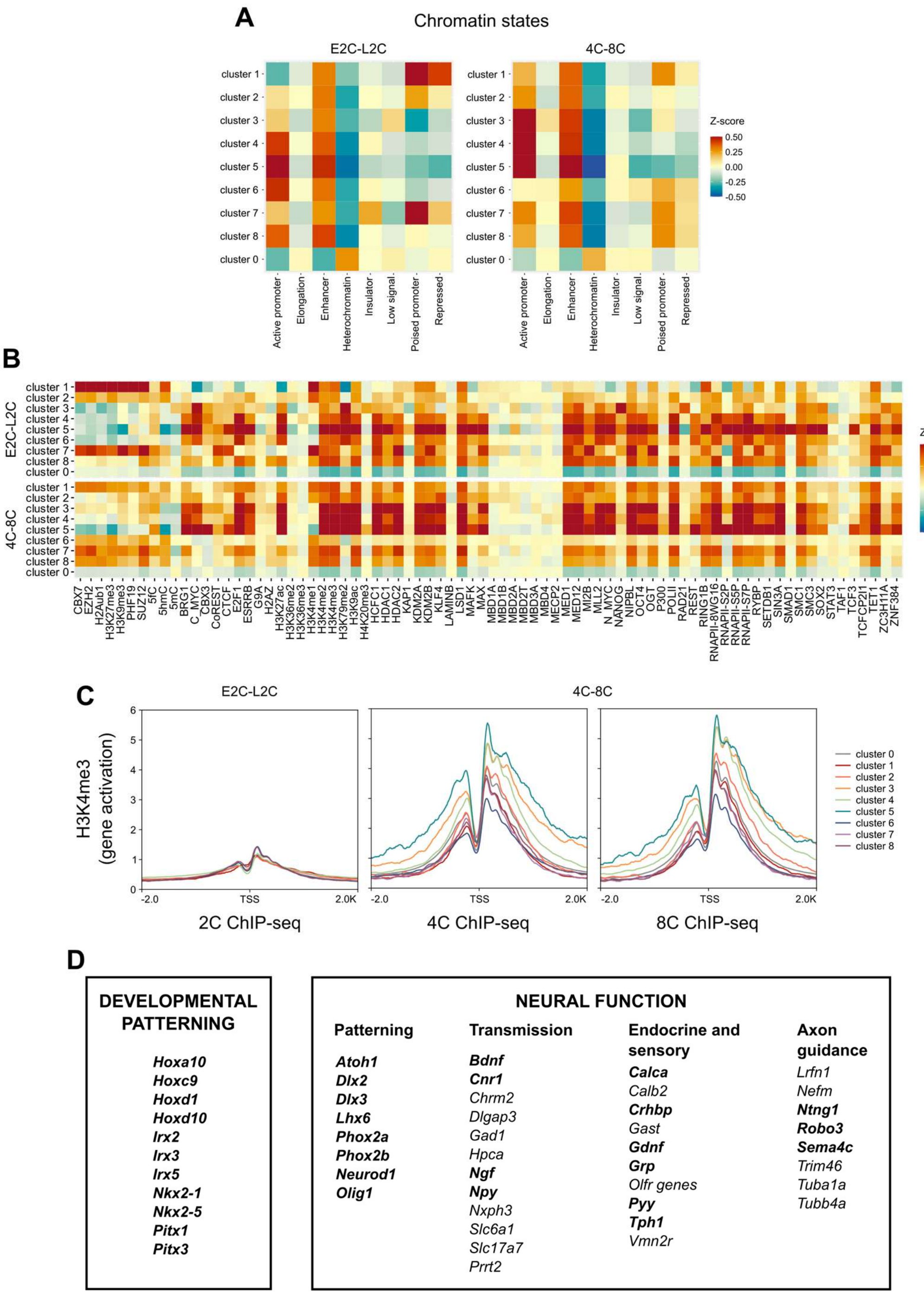


**Figure S5. Chromatin dysregulation in response to inhibition of DNA replication.**

**(A)** Heatmaps showing the degree of overlap (Z-score) of the TSSs of genes from each of the cluster (rows) from the E2C-to-L2C (left) and 4C-to-8C (right) time windows analyzed, with different chromatin states (columns). Color intensity indicates relative levels of overlap, with blue denoting lower and red higher overlap. Chromatin state classification was obtained from Juan et al. (2016).

**(B)** Heatmap showing the overlap (Z-score) genes from each of the cluster (rows) from the E2C-to-L2C (left) and 4C-to-8C (right) time windows analyzed, with individual histone modifications and chromatin binding factors (columns). Color intensity indicates relative levels of overlap, with blue denoting lower and red higher overlap. Data was obtained from Juan et al. (2016).

**(C)** Average profiles of H3K4me3 ChIP-seq signal on a 4 kilobase (kb) region surrounding the TSS of genes from each of the clusters defined in the E2C-to-L2C (left) and 4C-to-8C (middle and right) developmental windows (color code shown on the far right). Data for ChIP-seq generated in 2-cell (left), 4-cell (middle) and 8-cell (right) embryos was obtained from Li et al. (2025).

**(D)** Genes from cluster #1 of the E2C-to-L2C window, grouped by function. Polycomb targets are indicated in bold.
